# Biphasic bacterial community assembly predicted from generalized first principles of monoculture growth and inferred species interactions

**DOI:** 10.64898/2026.07.10.737752

**Authors:** Isaline Guex, Melanie L. Stäubli, Anna Sintsova, Vladimir Sentchilo, Senka Causevic, Adline Vouillamoz, Clara Bailey, Hans-Joachim Ruscheweyh, Shinichi Sunagawa, Christian Mazza, Jan Roelof van der Meer

## Abstract

Microbial communities occur in all habitats, yet how individual growth on available nutrients scales to community assembly remains poorly understood. This gap stems largely from the unknown effects of species interactions. These interactions arise because individual populations both consume and transform primary substrates into metabolites exploitable by others, and because parasitic and predatory mechanisms can release cellular building blocks that enable nutrient reuse. Here, we present a mathematical framework that predicts community growth and compositional succession from monoculture growth kinetics, resource availability, and species interaction parameters. To parametrize species interactions, we use a simulated-annealing optimization algorithm to search parameter space for sets that minimize the difference between modeled community growth and experimental time series from soil microcosms inoculated with defined communities of 20 or 21 soil isolates, with or without an opportunistic bacteriovorous member. The optimized interaction parameter sets were then used to predict growth dynamics in an independent 21-member community and in species drop-out communities. We find that community development is biphasic: an initial phase dominated by competition for primary resources driven by inherent strain growth kinetics, followed by a phase governed by cross-feeding and biomass formation on released byproducts. Paired metatranscriptomic analysis corroborated predicted shifts in individual growth states and revealed metabolic repurposing associated with the sudden renewed availability of metabolites and cellular building blocks. Model simulations that excluded species interactions reproduced only one-fifth of the observed community biomass, highlighting the importance of cross-feeding for soil community growth. Overall, models that integrate monoculture growth kinetics with inferred species interactions can predict the dynamics of medium-complexity communities from starting inocula even when environmental nutrient composition is largely unknown.

## INTRODUCTION

Growth is a core property of bacterial populations, determined by species-inherent metabolic capacities, biotic interactions, available resources, and the physicochemical properties of their habitat. The kinetics of single-strain cultures growing on finite resources are well studied: Monod’s original mathematical formulation of growth kinetics (*1*), refined many times since (*2-4*), is now widely accepted. By contrast, microbes in natural environments or host-associated communities rarely exist as pure cultures; they occur within diverse multispecies assemblages (*5, 6*). Because microbial communities are crucial for health and ecosystem functioning, a central question is how to extend single-species growth formalisms to describe growth of multiple species under defined substrate and habitat conditions (hereafter “community growth”).

The main challenge is that multispecies mixtures generate dynamic interactions that can enhance or inhibit individual development, and these interactions are not captured by the original Monod framework. Although several approaches have been proposed - including consumer-resource models (*7-13*), or generalized Lotka-Volterra formulations (*14-19*), none consistently link emerging species interactions to Monod-type, substrate-dependent growth kinetics for individual species. A useful community-growth framework therefore needs two components: (1) a description of substrate conversion to biomass that retains the classical dependence of growth rate and substrate availability, and (2) a way to represent how emergent interactions modify each species’ substrate-dependent growth kinetics.

We and others previously proposed a chemical-reaction-network formalism that is analogous to Monod growth kinetics, in which growth of a bacterial population on a single substrate arises from two coupled processes: direct consumption of the primary (added) substrate and later uptake of released metabolic byproducts (*20, 21*). In monocultures, measured biomass yield and growth rates reflect the combined effect of primary-substrate use and byproduct recycling, because the two contributions are difficult to separate experimentally. In multispecies communities, however, byproducts released by one species can be taken up by another, and vice-versa; this cross-feeding can be captured mathematically as species-specific waste production that serves as additional substrate for growth (*20, 21*). We previously showed that modeling two-species competition on a single resource is substantially improved by including such cross-feeding, compared with alternative community-growth models (*21*). Thus, accounting for metabolite exchange is essential for describing community dynamics – a conclusion supported by many experimental studies showing widespread metabolite release during growth (*22-24*), and that demonstrate the important role of byproduct-mediated cross-feeding in shaping community composition and diversity (*25-29*), stability (*25, 30*) and spatial structure (*31, 32*).

The major objective of the current work was to extend the cross-feeding growth model for cocultures (*21*) to community level including emerging species interactions (*Methods*). We hypothesized that two main components exist that underly species interactions. The first could be called substrate-mediated interactions (SMINT), such as being the result of cross-feeding on metabolites leaked during transformation of the primary available substrates in the system. SMINT can have a positive but also a negative component if the released metabolites are growth inhibitory to the other species. The second component we call parasitic interactions (PARINT), being the result of predation or damage inflicted by one species on the other, as a result of which cell contents leaks out and can be again used as growth substrates or biomass building blocks by others.

In absence of available interaction parameter sets, we use a simulated-annealing optimization (SAO) algorithm that iteratively searches parameter values to minimize the differences of model-predicted and experimental observations of community growth (Fig. 1). Communities consisted of 20 or 21 soil isolates inoculated into sterile soil microcosms as described previously (*33, 34*). Optimized SMINT- and PARINT-parameter sets were then used to predict growth and species succession in an independently started soil-inoculated 21-member community. Paired metatranscriptomic analysis was used underpin growth-related gene expression and explain metabolic differences among community members appearing as a result of species interactions. Finally, we used the model to predict outcomes for species drop-out communities and compared interaction patterns in soil-grown versus mixed liquid culture conditions for the same community.

**FIG 1.**
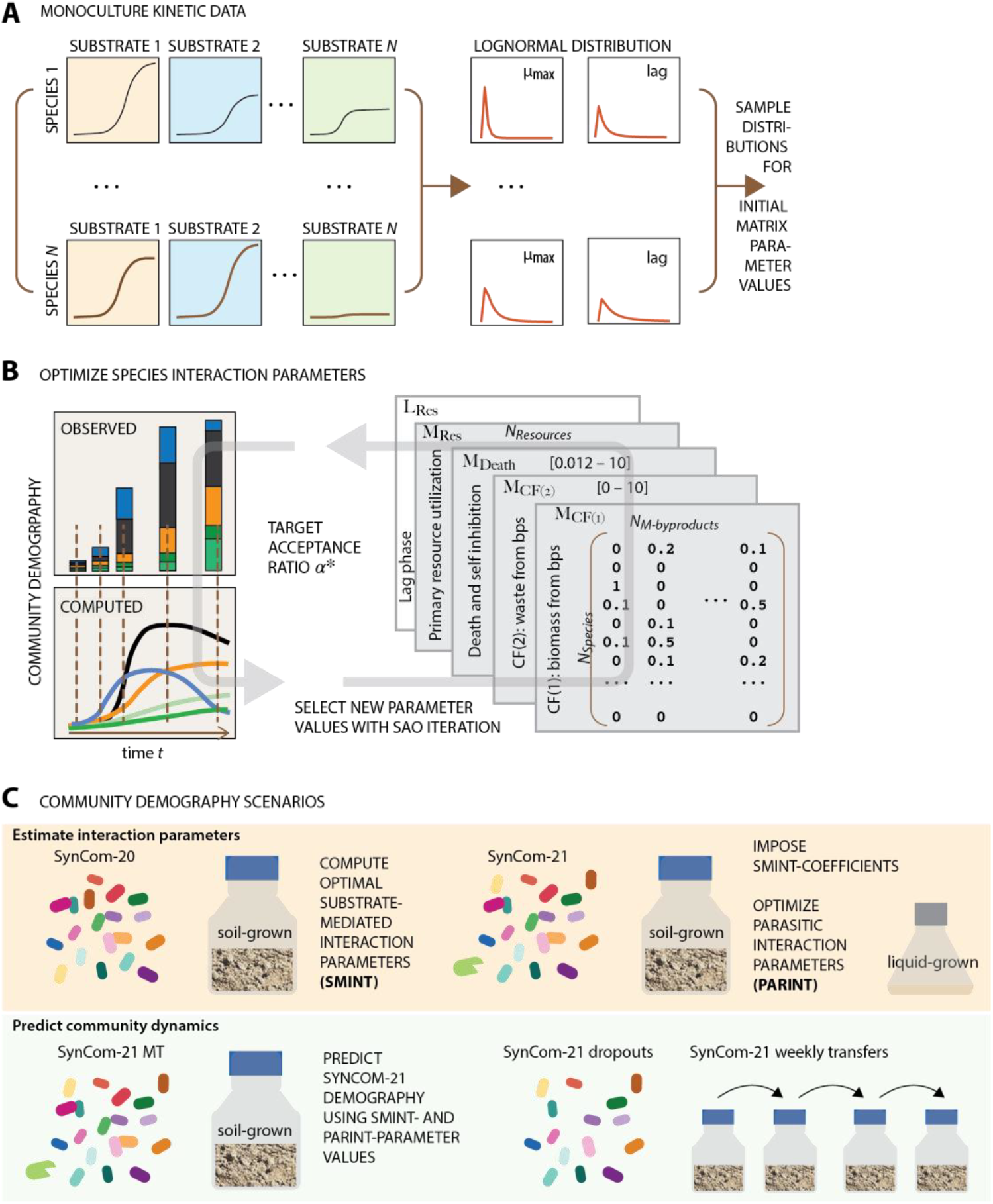
Concept and design of the community demographic model. **A)** Individual monoculture growth kinetics on different substrates are used to construct per-species log-normal distributions of maximum specific growth rates and lag times, and to initiate community growth simulations. **B)** Schematic representation of a simulated annealing optimization (SAO) round to estimate species interaction parameters. The SAO-algorithm searches parameter space of the interaction matrix values feeding into the demography model to narrow the differences between observed and simulated absolute species abundances until the target acceptance ratio is reached. See *Methods*. **C)** Soil-grown SynCom-20 data are used to estimate substrate-mediated interaction parameter values (SMINT) that are next imposed to SynCom-21 experiments to estimate parasitic interaction parameters (PARINT). The optimized SMINT- and PARINT-parameter sets are then used to predict community growth outcomes starting from new initial species abundances with initiated monoculture growth kinetic values in independently experimentally verified scenarios. MT, metatranscriptomics experiment.

## RESULTS

### A mathematical community growth model based on monoculture growth rates and inferred interspecific interactions

We generalize a community growth model based on first principles of Monod kinetics, under the assumption that community growth is an extension of inherent growth physiology of the individual bacterial strains that constitute the assembled community, constrained by the dynamic utilization of primary resources and widened by appearing metabolic byproducts released by other community members (Fig. 1, *Methods*). Growth rates under unknown nutrient conditions can be simulated by random sampling of growth rate distributions measured for individual strains under diverse substrate conditions (Fig. 1A). Utilization of byproducts for biomass formation is coded as being the result of metabolic waste production or of parasitic and predatory release and reuse (*Methods*, **eq. 1 &2**). The parameter values describing biomass formation from byproduct utilization are estimated by an SAO-algorithm that minimizes the discrepancy between observed and simulated absolute population abundances over time (Fig. 1B). Optimized parameter sets can subsequently be used for predictions of growth dynamics in unseen communities (Fig. 1C).

The substrate utilization capacities of each of the 21 soil isolates (table S1) were determined in pure culture from Biolog growth plates with 30 individual carbon substrates, complemented by growth assays in complex and defined media, and in aqueous extracts of soil organic matter. In addition, the total population size of each individual strain grown alone in sterilized soil microcosms was recorded. All strains grew on complex medium (PTYG) and soil extract, but to varying extents on individual substrates (Fig. 2A). Monoculture-observed maximum specific growth rates were represented by log-normal distributions with few substrates supporting high and most substrates resulting in low growth rates (Fig. 2B). Monoculture observed maximum population sizes after one week in soil microcosms differed up to 200-fold (Fig. 2C), which was unrelated to the width of the per-species log-normal growth rate distribution (compare, e.g., growth of *Microbacterium* or *Luteibacter* with *Pseudomonas* sp. strain 1), attesting to the varying capacity of the species to colonize the soil with its available resources. The simulated maximum population biomass after one week of monoculture growth in soil microcosms, using the measured kinetic parameters and their fitted log-normal distributions, was very similar to the observed population densities (Fig. 2C, biomass converted to cells per gram of soil), indicating that the growth model accurately captures observed monoculture data in absence of any species interactions.

**FIG 2.**
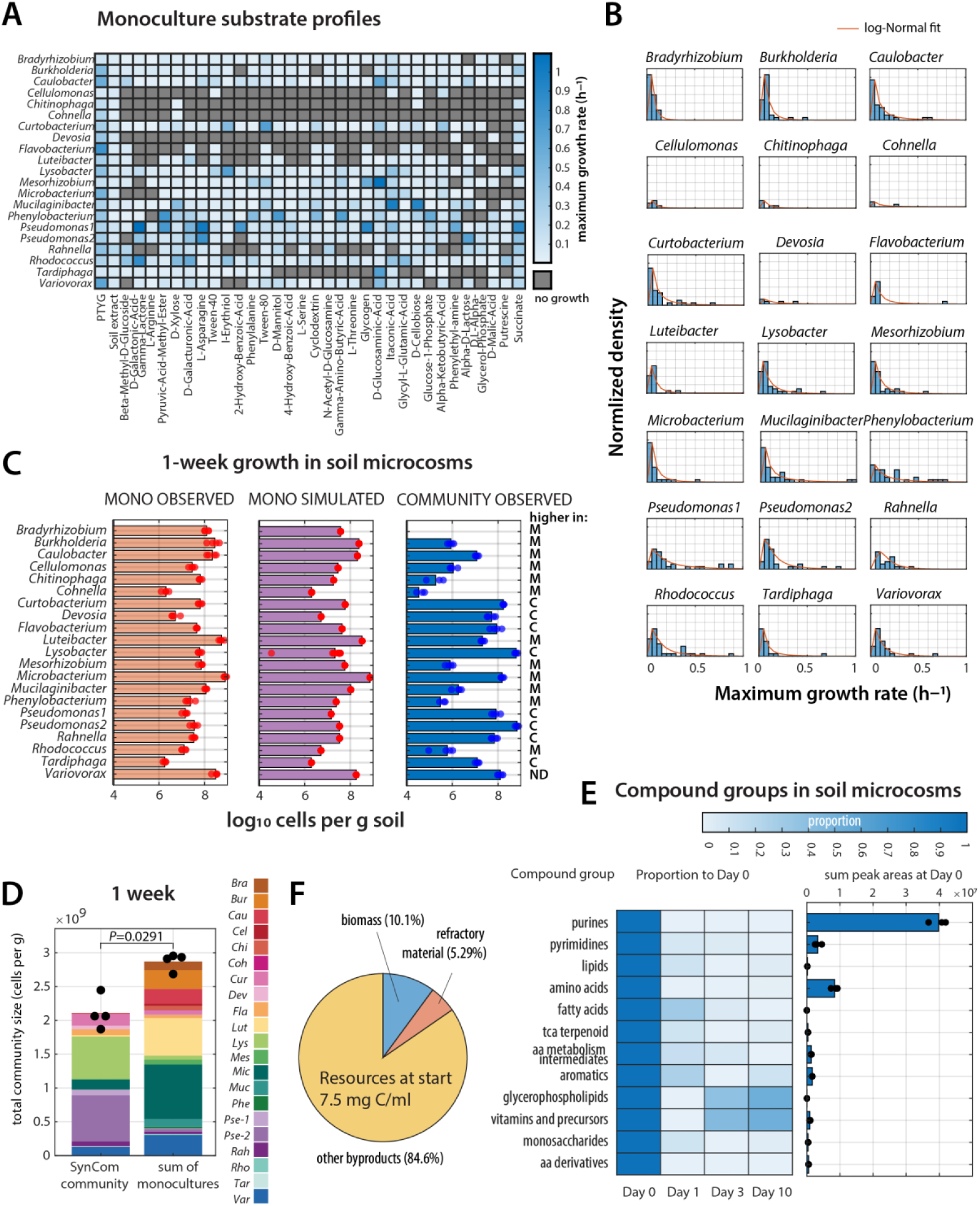
Monoculture growth kinetic profiles of soil isolates. **A)** Monoculture observed substrate utilization profiles (fitted maximum specific exponential growth rates, in h^−1^; as heatmap per color scale on the right), and **B)** deduced per-strain maximum growth rate profiles across all tested substrate conditions (normalized density plots with log-normal fitted density curves in red). Profiles of (B) are used for the initial random sampling of growth rates per substrate category in simulations. **C)** Comparison of soil-microcosm observed monoculture growth after 1 week (*n* = 4 replicates), of simulated monoculture growth using kinetic values of A and B, and observed individual species growth within the 21-member community in soil (*n* = 4 replicates). M, higher in monoculture; C, higher in community; ND, no difference (*P* < 0.05; two-tailed paired t-test with FDR correction). **D)** Mean total community size in soil microcosms (bars) after 1 week of growth (*n* = 4 replicates as individual dots, stacked by strain identity) compared to the mean of total sums of monoculture-colonized soil microcosms (arbitrarily grouped to *n* = 4 replicates). *P* value from two-sided Wilcoxon ranksum test. **E)** Measured compounds (targeted HILIC-MS) in aqueous extract of soil microcosms at start (non-inoculated, Day 0), and over time of SynCom-21 growth (Day 1, 3 and 10). Compounds grouped per category on the left; amounts represented as the mean sum of peak areas (bar diagram on right, *n* = 3 replicates as individual dots), and as proportion of Day 0 per category (heatmap, colorscale on top). **F)** Estimated carbon resource balance in SynCom-21-grown soil microcosms at day 10: total measured organic carbon (resource at start), bacterial biomass (converted from cell numbers with estimated per-species dry weights), refractory material (fraction of quantifiable compound in HILIC-MS at Day 10 compared to Day 0, table S2), and the *other* byproducts (difference of total resources at start, biomass and refractory material).

Compared to growth within an inoculated community in the soil microcosms, about 60% of the strains (13 out of 21) performed better in isolation and the others better in community (Fig. 2C). In contrast, none of the species in monoculture microcosms reached a population size equivalent to the total observed community size, although (an arbitrary) sum of all monoculture sizes slightly surpasses the total community size (Fig. 2D, *P* = 0.0291, two-sided Wilcoxon ranksum test). Assuming the same resource availabilities in the starting soil material for the microcosms, this indicates that species growing in the community mobilize the available nutrients in a different manner than any species alone, for example, as a result of species interactions. This is evident for strains with higher maximum biomass in community than in isolation (e.g., *Lysobacter* and *Pseudomonas* sp. strain 2; Fig. 2C).

Aqueous-extractable compounds from the soil microcosms categorized into 12 chemical groups by hydrophilic interaction liquid chromatography coupled to mass-spectrometry (HILIC-MS) analysis were dominated by purines, amino acids and pyrimidines, suggesting these to be the primary available substrates for growth (table S2). Substrate levels decreased rapidly in growing communities in the first 1– 3 days, with little refractory compound accumulating at later time points (e.g., categories phosphoglycolipids and vitamins and precursors; Fig. 2E, table S2). In quantitative terms, an estimated 10.1 % of C mass of the measured primary resources was converted into bacterial biomass, 5.3 % was detected in the aqueous phase after one week, and the rest is C waste (Fig. 2F; refractory non-water extracted compounds or disappeared from the system as e.g., CO_2_).

### Substrate-mediated interactions are major enablers for community growth

Next, we sought to dissect the contributions of observed community growth in soil microcosms attributable to either inherent individual strain kinetic differences or to emerging species interactions. We first focused on interactions arising from metabolite cross-feeding from primary substrate conversion (SMINT). For this we used a 20-member defined representative soil community (*33, 34*) (SynCom-20; no inclusion of known predators such as *Lysobacter*; table S1). The SynCom-20 grown in soil microcosms rapidly expanded in total size in the first 3 days, reaching its maximum total biomass after day 7 followed by a slower decline (Fig. 3A, biomass converted from measured cell numbers and average dry weights per strain, see *Methods*; table S1). The community became dominated by 5–6 strains, which showed different succession patterns. Starting from even initial strain proportions, the relative abundances of *Pseudomonas* sp. strains 1 and 2 and *Rahnella* increased after one day, followed by a steep increase in *Flavobacterium* and a more gradual increase in *Caulobacter, Variovorax*, and other minor species (Fig. 3A).

**FIG 3.**
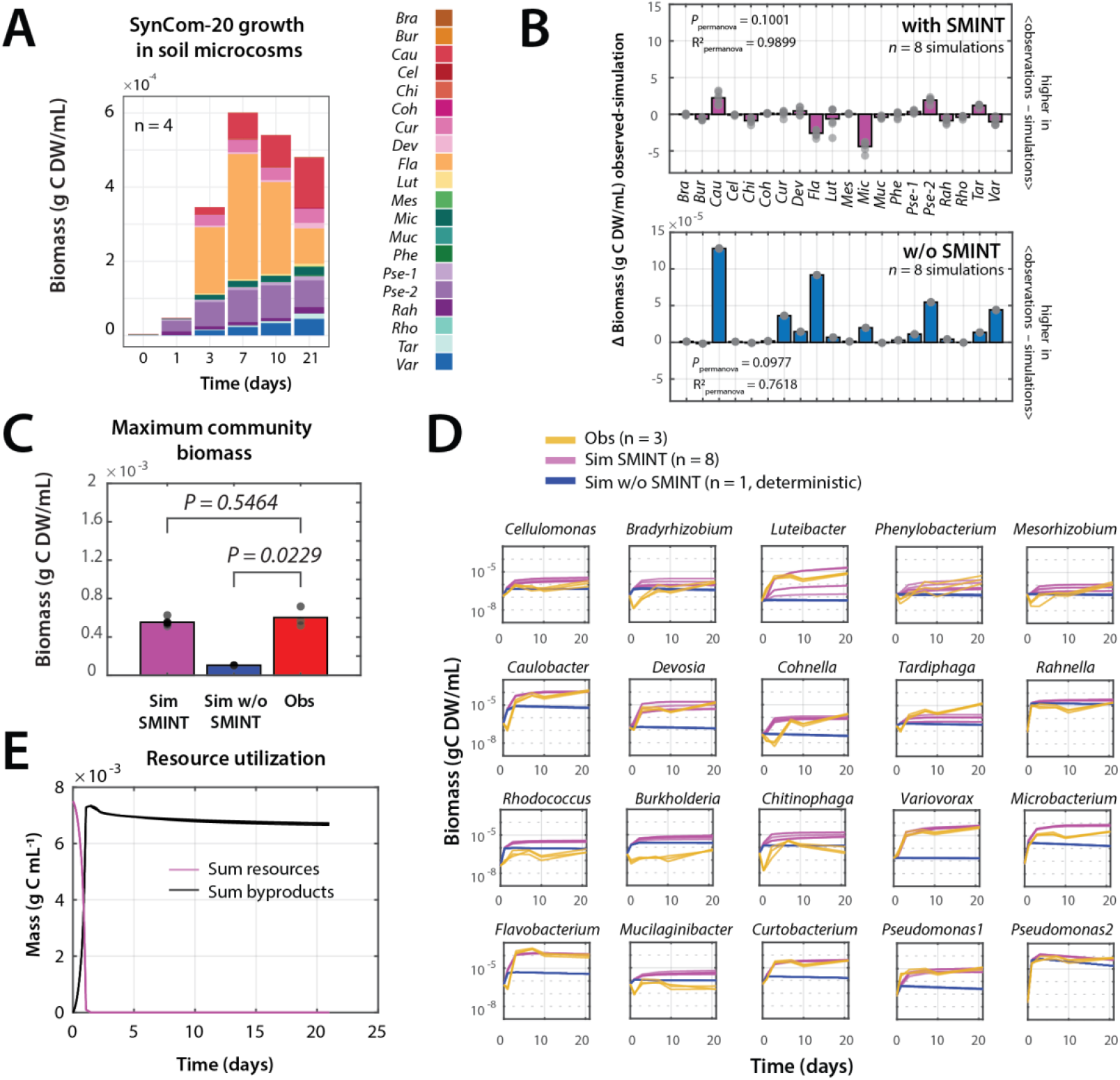
Estimation of substrate-mediated interactions during SynCom-20 community growth. **A)** Mean absolute growth of the SynCom-20 community in soil microcosms (*n* = 4, expressed as biomass in g C dry weight (DW) ml^−1^ soil water; stacked by absolute strain population sizes as per color scale). **B)** Mean per species biomass difference (bars) at day 21 between observed soil microcosm-growth and community growth simulations in presence of substrate-mediated interactions (with SMINT) or in their absence (w/o SMINT, *n* = 8 simulations shown by individual dots). *P* values derived from PERMANOVA using Bray–Curtis distances. R^2^ represents the proportion of the total explained variance by either observation or simulation. **C)** Total maximum community biomass growth for simulations (Sim) with (*n* = 8), or without SMINT (*n* = 1, since deterministic), compared to experimental data (Obs, *n* = 3). *P* values obtained from a permutation test of Cliff’s delta index, with n = 100,000 permutations. **D)** Individual observed (Obs, yellow) and simulated strain growth in the SynCom-20, in absence (blue) or in presence of SMINT (magenta). **E)** Simulated summed resource utilization and byproduct formation with optimized SMINT-values (*n* =8 simulations).

SMINT-parameters were estimated by the SAO-algorithm (*Methods*) trained on 5 out of 8 combined replicate datasets from two independent soil-grown SynCom-20 experiments, with 3 random picked replicates left out for subsequent validation. SAO-estimations were repeated 10 times from growth kinetic values picked randomly from the log-normal distributions (Fig. 3B). Community growth simulations with optimized SMINT-parameters reproduced the observed community growth quite well, both in terms of individual population sizes, trajectories and final total community biomass (Fig. 3B-D; mean theta cosine trajectory value of 0.4445, Table 1). Observed growth trajectories of some strains showed an initial temporary decrease in population size (e.g., *Cohnella, Rhodococcus, Burkholderia*; Fig. 3D), which may be due to a decrease of cell viability with respect to the measured inoculum size by flow cytometry. Because growth simulations are initiated with the measured inoculum size, an actual loss of inoculum viability will lead to an overestimation of simulated population growth. The 10 independent SAO-simulations started from freshly initiated subsampled kinetics showed good parameter convergence and little variation among replicates (Fig. 3D). In contrast, drawing random kinetic starting values for SMINT-parameters from the respective species’ growth rate distributions without further SAO-runs resulted in individual population growth strongly deviating from the experimental observations, despite good convergence of community trajectories (fig. S1, Table 1). This suggests that the SAO-algorithm is not overfitting SMINT parameters, but capturing relevant biological meaning.

**Table 1.** Theta cosine similarity between observed and simulated community trajectories.

| Experiment |  |  | With SMINT and/or PARINT<sup>a</sup> |  | No SMINT- or PARINT-parameters |  |
| --- | --- | --- | --- | --- | --- | --- |
| SynCom | Habitat | Type<sup>a</sup> | Theta<sup>b</sup> | P value<sup>c</sup> | Theta<sup>b</sup> | P value<sup>c</sup> |
| SynCom-21 | Soil microcosms | SAO-SMINT/PARINT | 0.7026 | 3.6×10<sup>-3***d</sup> | 5.77×10<sup>-2</sup> | 0.235 |
|  |  | SAO-SMINT | 0.3473 | 4.1×10<sup>-3**</sup> | 4.76×10<sup>-2</sup> | 0.3815 |
| SynCom-21 *MT* | Soil microcosms - metatranscriptomics | Prediction | 0.6749 | 2.0×10<sup>-4***</sup> | 0.3052 | 1.1×10<sup>-2**</sup> |
| SynCom-21 *L* | Liquid mixed culture | SAO-SMINT/PARINT | 0.9485 | 7.0×10<sup>-3***</sup> | 0.2389 | 7.6×10<sup>-3</sup> |
| SynCom20 | Soil microcosms | SAO-SMINT | 0.4445 | 1.4×10<sup>-2**</sup> | 0.2499 | 8.8×10<sup>-2*</sup> |
|  |  | SAO-Random SMINT | 0.223 | 2.1×10<sup>-2**</sup> | 0.2376 | 8.5×10<sup>-2*</sup> |
| SubCom C1 | Soil microcosms | Prediction | 0.335 | 0.123 | 0.2276 | 0.2038 |
| SubCom C2 | Soil microcosms | Prediction | 0.3252 | 0.1288 | 0.2137 | 0.2513 |
| SubCom C3 | Soil microcosms | Prediction | 0.5929 | 4.0×10<sup>-2**</sup> | 0.5828 | 4.1×10<sup>-2**</sup> |

a) SAO, simulated annealing optimization algorithm. SMINT, substrate-mediated interactions; PARINT, byproduct usage from parasitic interactions and cell lysis; random, random starting kinetic parameter values. Prediction, direct use of the SynCom-21 SAO-SMINT/PARINT-parameter values without further SAO-searches.
b) Theta cosine similarity between community observations and simulations, including all species, time points and replicates. Value > 0.7 indicates high similarity in community trajectories; 0.3 – 0.7: moderate similarity; tendency to develop in the same direction; 0 – 0.3 weak similarity and tendency to follow the same trajectories.
c) P value represents the probability that permutations of the simulated time series values would produce a similarity at least as extreme as the observed one, tested against the null hypothesis that the temporal order of the community trajectory does not carry meaningful information about its similarity with the reference and, therefore, random permutations of the temporal values would produce cosine similarities comparable to the observed.
d) Results of the test remain exploratory due to the small number of experimental replicates ( $n = 3 - 5$ ). \*\*\*, $P < 0.01$ indicates strong evidence against the null hypothesis; \*\* $P < 0.05$ , indicates evidence against the null hypothesis; \* $P < 0.1$ , suggests potential relationship.

The effect of including SMINT on community growth was then simulated by setting the SMINT-parameter values to zero. In this case the strains can only utilize and compete for primary available resources, which is determined by their individual kinetic parameters (i.e. maximum growth rates, yields and half-rate constant K_S_). Simulated SynCom-20 growth without SMINT-effects starkly underestimated the experimentally observed population sizes of all strains, most evident for the dominating members (Fig. 3B). Predicted growth trajectories had overall lower scores (cosine theta value of 0.2499, Table 1). In addition, the total estimated SynCom-20 biomass in simulations without SMINT only reached one-fifth of the experimentally observed values (Fig. 3C, *P* = 0.0229, permutation test on Cliff’s index). This large difference would indicate that substrate-mediated metabolite cross-feeding is a key component for productivity of soil communities and benefits community member growth widely.

Substrate utilization as a function of community growth was also correctly reproduced in the SMINT SynCom-20 simulations (Fig. 3E), mirroring the rapid decrease in measured extractable soil compound concentrations within the first day (Fig. 2E). Different simulation runs indicated fast per-strain byproduct appearance, a decrease between day 3 and 5, and then a slow stabilization at around 6.5 g C dry weight (DW) per ml (Fig. 3E). This suggests that the decrease in primary (water-extractable) resources is the consequence of the rapid consumption and increase in the population sizes of fast-growing community members, followed by a second phase of slower community growth when waste products dominate and are being consumed (Fig. 3D).

### Estimating parasitic interactions from predation and cell building blocks turnover

In a second step we extended the community growth model with parasitic species interactions (PARINT) resulting from predation and the corresponding reutilization of cellular building blocks. For this we used time series of soil microcosm-grown communities of 21 strains (SynCom-21) that now included the opportunistic bacterial predator *Lysobacter* (table S1). We assumed that we could keep the previous optimized SMINT-parameter sets, since the same 20 soil bacteria would still be present in the SynCom-21 community, and use the SAO-algorithm to search only additional SMINT-values necessary for inclusion of *Lysobacter* and estimate PARINT-parameter sets that we expected would be the result of *Lysobacter* opportunistic predatory behaviour. Similar as for the SynCom-20 optimization, the SAO-iterations explored parameter sets that would minimize the discrepancy between (new) experimental data of SynCom-21 community growth and the growth dynamic simulations.

Including *Lysobacter* in the SynCom inoculum considerably changed the community dynamics (Fig. 4A with SynCom-21 vs Fig. 3A with SynCom-20). Total community size again increased mostly during the first three days followed by a decline, but the dominant species were different in the SynCom-21, with notable higher proportions of *Lysobacter* and *Pseudomonas sp*. strain 2 and lower proportions of *Caulobacter, Variovorax, Flavobacterium* and *Microbacterium* (Fig. 4B, PERMANOVA *P* = 0.0278).

**FIG 4.**
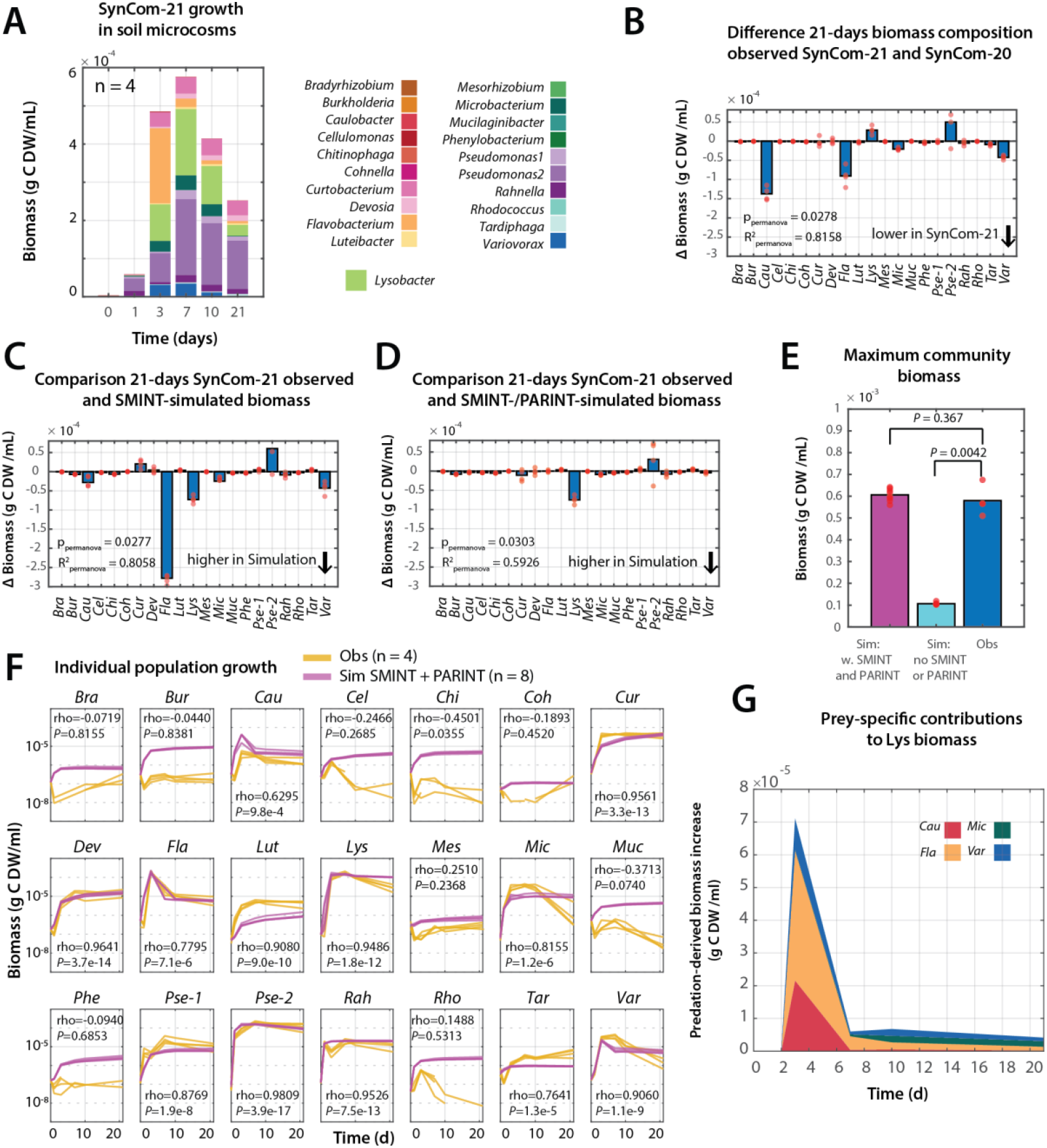
Estimation of parasitic interactions on growth of a 21-member defined community with opportunistic bacterial predator. A) Mean absolute growth of the SynCom-21 community including the opportunistic predatory bacterium *Lysobacter* in soil microcosms (*n* = 4, expressed as biomass in g C DW ml^−1^ soil water; stacked by absolute individual community member population sizes as per color scale). B) Mean per species biomass difference (bars) at day 21 between observed soil microcosm-grown SynCom-20 and SynCom-21. C) and D) As (B) but for the differences between the experimental SynCom-21 data and growth simulations with optimized SMINT-parameters only (i.e., no parasitic release of cell building blocks), or with SMINT- and optimized PARINT-parameters (*n* = 8 simulations shown by individual dots). *P* values and R^2^ from PERMANOVA using Bray–Curtis distances as in Fig. 3. E) Total maximum simulated (sim, *n* = 8) SynCom-21 community biomass growth with or without SMINT- and PARINT-parameters, compared to experimental data (Obs, *n* = 4). *P* values from a permutation test on the Cliff’s delta index with n = 100,000 permutations. F) Individual observed (Obs, yellow) or simulated SynCom-21 species growth with both SMINT and PARINT (Sim, magenta). G) Estimated prey-specific *Lysobacter* biomass gain over time from predation (species colors as in legend).

The importance of including PARINT-effects in SynCom-21 community demography can be deduced from the difference of simulations with only the SMINT- or with both SMINT- and PARINT-parameters. Using only SMINT-values overestimated biomass growth of *Flavobacterium, Variovorax* and *Caulobacter* in the SynCom-21 (Fig. 4C), whereas inclusion of PARINT-parameters improved the simulated SynCom-21 population abundances for all species in comparison to the experimental data (Fig. 4D, Table 1). The model was not completely correct for *Lysobacter* (overestimated) and *Pseudomonas* sp. strain 2 (underestimated; Fig. 4D). Setting both SMINT- and PARINT-parameters to zero reduced the simulated SynCom-21 biomass to only approx. 20% of the experimentally observed levels (Fig. 4E, *P* = 0.0042, permutation test on Cliff’s index), again indicating the crucial importance of species interactions for community productivity. Individual population growth was correctly recapitulated by the SMINT/PARINT-simulations, which even captured an observed decrease in abundances of *Flavobacterium, Caulobacter* and *Variovorax* after 3 days (Fig. 4F). In contrast, growth was overestimated for some of the less abundant species (e.g., *Phenylobacterium* or *Bradyrhizobium*, Fig. 4F), possibly again because their actual viability in the soil was lower than assumed from their inoculum size.

To deduce the most likely prey for *Lysobacter*, we identified SynCom-strains grown in soil microcosms that showed a large biomass reduction in presence of *Lysobacter* (Fig. 4B); further corroborated by simulations with and without PARINT parameters (Fig. 4C &D). Four species met this criterion: *Flavobacterium, Caulobacter, Microbacterium* and *Variovorax* with more than 92% biomass reduction in presence of *Lysobacter*. Estimations from the PARINT-parameter values suggested that *Lysobacter* biomass growth from predation is biphasic, with a first strong peak between day 2 and 5, and a second minor contribution thereafter (Fig. 4G).

### Asymmetrical species interactions underly community biomass formation from byproducts

The combined SMINT- and PARINT-effects can be summarized as the amount of per-species biomass formed from byproducts being released by others and over time. *Pseudomonas* sp. strain 2 is the first to grow in both SynCom-20 and SynCom-21, and the simulations indicate that the population of this strain alone would be responsible for byproducts in the range of 2×10^−3^ g C ml^−1^ (Fig. 5A &B). Byproduct formation coincides with growth as would be expected for release of metabolic intermediates and halts when the population reaches stationary phase (Fig. 5C). Seen from the perspective of biomass formation, *Pseudomonas* sp. strain 2 also consumes byproducts released by other members in the early stages of SynCom-20 community growth (12 and 24 h), whereas the cross-feeding on byproducts from *Pseudomonas* sp. strain 2 emerges in later stages of community growth (48 h and beyond, Fig. 5D). The byproduct utilization matrix also suggests how other abundant strains in the SynCom-20 could profit from dynamic byproduct release (e.g., *Flavobacterium, Caulobacter, Variovorax* and *Microbacterium*, Fig. 5D). By contrast, simulations suggest that *Devosia* in SynCom-20 is a net producer but a poor consumer (Fig. 5D). In the presence of *Lysobacter* in SynCom-21, the byproduct utilization matrix changes (Fig. 5E), suggesting more production and utilization of byproducts by *Pseudomonas* sp. strain 2 in the first stages (12 and 24 h), and a change to *Flavobacterium* as producer with *Lysobacter* becoming a consumer (from 48 h onwards – consistent with their prey-predator relation). At later stages (168 h), *Lysobacter* is changing into a byproduct producer, consistent with the idea of byproducts from cell lysis building blocks being turned over, leading to new byproducts from which other species profit (Fig. 5E, see *Methods* for the modeling concept).

**FIG 5.**
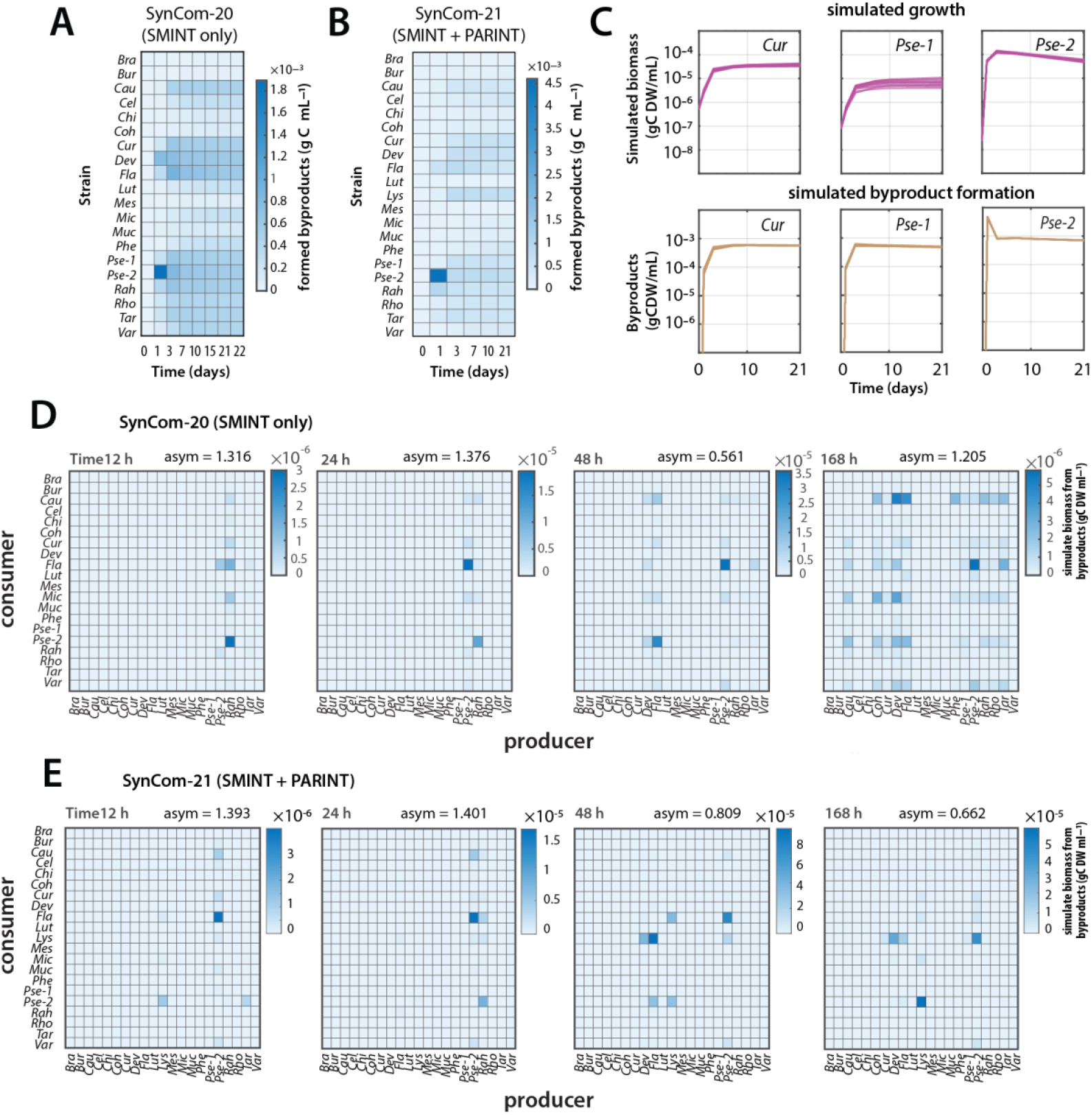
Estimated interaction partners and growth effects from byproduct usage. **A)** and **B)** Mean formed byproducts (in g C per ml) from the individual strains over time in the simulated SynCom-20 (with SMINT-parameters) or SynCom-21 communities (including both SMINT and PARINT). **C)** Example of the dynamics of byproduct formation for three strains in simulated SynCom-21 growth. **D)** and **E)** Interaction matrices of byproduct producers versus consumers at specific time points during SynCom-20 (only SMINT) or SynCom-21 simulated growth (SMINT and PARINT). asym, measure for the symmetry of the matrix (0 corresponding to completely symmetrical, a value of 2 to maximal asymmetry).

### Simulation predictions of dynamic interactions in a SynCom-21 community

So far, the simulation models with optimized SMINT- and PARINT-parameter values explained and reproduced the observed growth of the SynCom-20 and -21 experimental datasets (Table 1). However, as these models search parameter space to match experimental results it may come as no surprise that their ‘fitting’ is relatively good. To gain more confidence in the biological significance of the optimized species interaction parameters, we next used the optimized SMINT- and PARINT-parameter sets to predict growth dynamics of an independently conducted SynCom-21 experiment starting with a different inoculum. For this additional SynCom-21 experiment (SynCom-21 *MT*, Table 1), we also generated paired time series metatranscriptomic and metagenomic data, which was used to infer *in situ* growth and to inform potential cross-feeding behavior from gene expression changes of uptake systems and metabolic pathways. In situ growth prediction was based on two tools: mTRAc and CoPTR. mTRAC is a binary growth state classifier, which uses proportional abundances in a set of single-copy marker genes to predict probabilities of growth ranging from 0 to 1, and classifies strains to be either in a growing or a non-growing state (*35*). CoPTR uses log_2_-ratios of highest-to-lowest (Peak-to-Trough) metagenomic sequencing coverage as a proxy for *in situ* growth rates (*36*).

Similar as in the previous experiments, growth of the SynCom-21 showed a succession of dominant populations, notably again *Pseudomonas* sp. strain 2, *Lysobacter, Rahnella, Flavobacterium* and *Variovorax*, as well as less dominant members (fig. S2A &B). The probabilities of growth, predicted by mTRAc for eight of the major abundant SynCom-21 members, corresponded quite well to the population dynamics predicted by the model that included SMINT- and PARINT-parameters (Fig. 6A, *n* = 10 simulations with randomly drawn parameter sets from the total of 40). All major community members were classified to be growing during early time points (12-38 h), but their probabilities of growth decreased at intermediate time points (38-168 h) in accordance with depleted primary nutrients (Fig. 2E). A sharp decline in growth probability for several strains (e.g., *Caulobacter, Flavobacterium* and *Variovorax*, Fig. 6A, fig. S2A &D) coincided with predicted and observed decreases in population sizes, which is an indication for predatory cell lysis by *Lysobacter* (Fig. 6A; compare to the previous SynCom-21 experiment in Fig. 4B). The growth dynamic behaviour of other strains, notably *Pseudomonas* sp. strain 2 and *Rahnella*, was more variable. The community demography model predicted for both a biphasic growth, the transition time of which corresponded exactly to a transient decrease in the predicted probability of growth from metatranscriptomic data at time point 24 h (Fig. 6A and higher time resolution in fig. S2F). At later time points (> 100 h) a larger variation in predicted probabilities of growth was observed for all strains (Fig. 6A), which probably reflects heterogeneity in population behavior from stationary phase cells, partial cell death, and renewed growth on released nutrients. Metagenomics-based predicted growth rates using CoPTR recapitulated similar population growth dynamics (i.e., steep decrease in predicted *in situ* growth rates for the 38-168 h time points) for fast-growing strains but failed to predict a decrease in *in situ* growth rates of lowly abundant strains (fig. S3).

**FIG 6.**
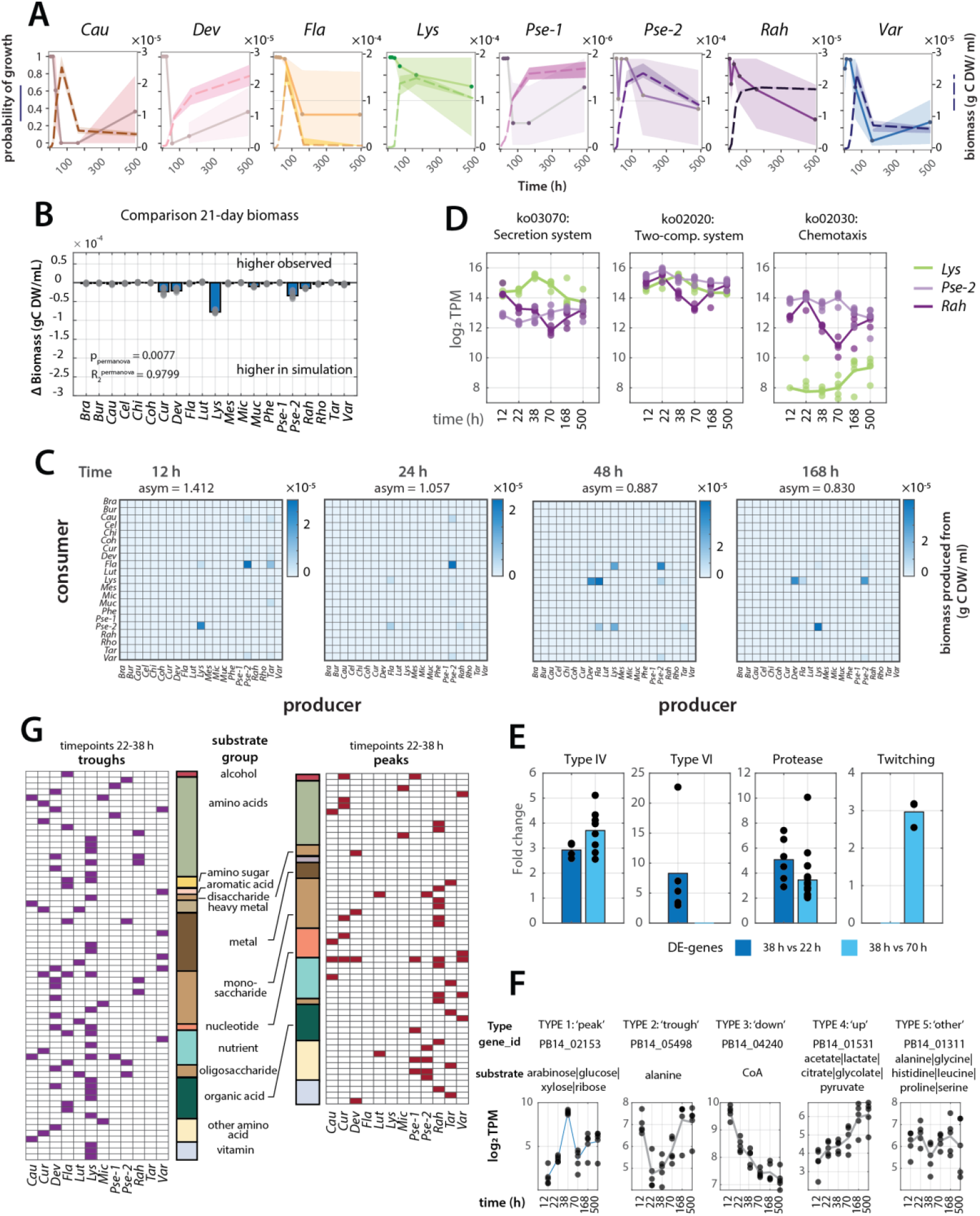
Predicted growth and interactions of an independently inoculated 21-member defined community in soil. **A)** Probability of growth for 8 of the 21 SynCom members estimated from expression of conserved housekeeping genes in metatranscriptomics (solid lines, scale on left axis), and predicted corresponding population growth from the demography model including both SMINT- and PARINT-parameters (dotted lines, scale on right axis. Note the change in axis scales between panels for optimal presentation). Shaded areas encompass the 25^th^ and 75^th^ percentiles of the replicate variation (*n* = 5 for observations, *n* = 10 for simulations). **B)** Comparison of the mean predicted and observed SynCom-21 biomass at day 21 per individual strain. Bars show the mean difference with dots the individual values. PERMANOVA analysis as in Fig. 3. **C)** Predicted species interactions as amount of formed biomass per individual population (consumer) as a function of released byproduct (producer) and at different time points during community development. **D)** Changes in summed expression levels (as log_2_ TPM) of KEGG-orthology pathways for three abundant populations in the SynCom-21, measured from metatranscriptomics. **E)** Mean fold-change (bars) of *Lysobacter* differentially expressed genes (DE, false-discovery rate < 0.05) at time point 38 h relative to either 22 h (dark blue) or 70 h (light blue) grouped by annotation key words relevant for its predatory life-style (dots are values from individual genes within key word groups). **F)** Categorization of time expression profiles of genes for specific uptake systems, two of which (TYPE 1 and 2) are reported in **(G)** with their predicted substrate groups and per genome (only shown when meta-transcriptomic data were reliable, fig. S4).

The predicted biomass at day 21 for each of the strains was in agreement with the experimental data, except for again overestimating *Lysobacter*’s biomass (Fig. 6B), whereas the total community biomass was 1.29-fold higher in the predictions than observed (fig. S2C, *P* = 0.0047, permutation test). The trajectories of both predicted and observed community dynamics aligned well (Table 1, cosine theta-score = 0.6749; fig. S2D), indicating that community growth was correctly predicted by the demography model.

The model further predicted that early species interactions would be dominated by *Flavobacterium* profiting from byproducts from *Pseudomonas* sp. strain 2 and *Pseudomonas* sp. strain 2 profiting from *Lysobacter*, then transitioning to include *Caulobacter, Lysobacter, Variovorax* and *Devosia* as consumers (Fig. 6C), with *Pseudomonas* sp. strain 2 as the overall largest byproduct producer for the community (fig. S2E). To support or refute these predictions, we analyzed the paired SynCom-21 *MT* metatranscriptomics dataset, focusing on three aspects: (i) expression of genes that could support the predatory behaviour of *Lysobacter*, (ii) altered expression of transporter genes and (iii) of KEGG-orthology (ko) pathways that may reflect strain adaptation to changing resources. Expression changes in genes associated with the ko-term *Bacterial secretion systems* (i.e., ko03070) at time points 38 and 70 h were 4-8 times higher for *Lysobacter* than for *Pseudomonas* sp. strain 2 or *Rahnella*, whereas those for *two-component systems* or *chemotaxis* were not (Fig. 6D). Statistically significantly increased *Lysobacter* expression at time point 38 h compared to either 22 h or 70 h also included type IV and type VI secretion system genes, genes annotated with ‘protease’ or ‘peptidase’, or ‘twitching motility’ (Fig. 6E), which are hallmarks of its predatory behavior (*37, 38*).

To detect strain-specific altered expression of transporter genes, we categorized their expression time profiles in five subtypes (e.g., TYPE 1: temporary increase between 22 and 38 h; TYPE 2: temporary decrease, Fig. 6F) among the strains with sufficient metatranscriptomic sequencing depth (i.e., at least 3 out of 5 replicates and 4 of 7 time points having reliable TPM-values, fig. S4A & B). Overall, most transporter genes showed increased expression over time (TYPE 4: ‘up’; Fig. 6F, fig S4B), indicating the strains to scavenge for scarcer nutrients. On average, between 4 and 8% of annotated transporters showed peaked expression (TYPE 1), and, for a subset of strains, 20-26% showing transient decreased expression (TYPE 2, fig. S4A &B). This distinction in TYPE 1 and 2 transporter expression broadly divided the behaviour of the SynCom-21 *MT* strains into two groups, with both *Pseudomonas* strains, *Rahnella* and *Microbacterium* in the category showing more transporters with peaked and fewer with transiently decreased expression at 22 h and 38 h, and the others (*Caulobacter, Devosia, Flavobacterium, Lysobacter* and *Variovorax*) with the opposite trend (i.e., fewer peaked and more decreased transporter expression; Fig. 6G, fig. S4A &B). Annotated transporter substrates covered a wide range of groups (Fig. 6G), including mono-and disaccharides, amino acids, organic acids, nutrients (e.g., N, P) and some vitamins (fig. S5). Both the peaked and the diminished transporter expression at 22 and 38 h may signal a sudden increase in available substrates, with peaks corresponding to those transporters that are under concentration feedback control (i.e., higher concentration leading to higher expression), and troughs to those that are insensitive (i.e., higher concentration outside meaning less expression needed to satisfy cellular needs). This sign for transitory increased substrate availability during community growth would then be the result of byproduct release from fast-growing dominant populations such as *Pseudomonas* sp. strain 2 and/or from lysed cells by *Lysobacter* predation.

### Species-dependent repurposing of metabolic pathway expression during community growth

To further corroborate this idea of temporarily increased availability of substrates in form of released metabolites, we compared the expression time profiles for genes within ko-pathways among the eight strains with sufficient metatranscriptomes sequence depth (fig. S4A & B). Principal component analysis separated the strains in similar groups as for the transporter gene expression, with both *Pseudomonas* strains and *Rahnella* displaying closer resemblance of ko-term expression patterns across time points, and the others showing wider trajectories (fig. S6).

According to dynamic expression classification, 11 ko-term expression types could be distinguished (e.g., transient, monotonous in- or decrease; fig. S4C). The per-strain attribution of ko-terms to those 11 categories were notably different. For example, *Caulobacter* and *Variovorax* showed a clear dominance of ko-terms attributed to category 9 (low expression at start, then sharp increase between time points 38 and 70 h, followed by a stable expression), *Flavobacterium* ko-terms were dominated by relatively high expression in early but lower in later time points, whereas *Devosia* and *Pseudomonas* sp. strain 2 showed a more or less equal attribution of ko-terms to all expression categories (Fig. 7A, fig. S4D). Despite this strong low-high transition in *Caulobacter* and *Variovorax* for almost all pathways, ko-terms for nucleotide metabolism, citric acid cycle and energy metabolism showed opposite trends (Fig. 7A). This suggests that cells remain active but take up available compounds feeding into those pathways, such as sugars and amino acids, to avoid *de novo* synthesis. Ko-terms in several strains (i.e., *Flavobacterium, Lysobacter, Pseudomonas* sp. strain 1 and *Rahnella*) showed remarkable lowest expression at time point 70 h, after which they slowly increased again. This is also indicative for temporary increased availablility of nutrients like pyruvate, succinate, certain amino acids, or nucleotides, which can be taken up and avoids the necessity of their biosynthesis (Fig. 7A). The sudden renewed nutrient availability is most likely the result of lysis of *Flavobacterium* by *Lysobacter* (as in Fig. 4G).

**FIG 7.**
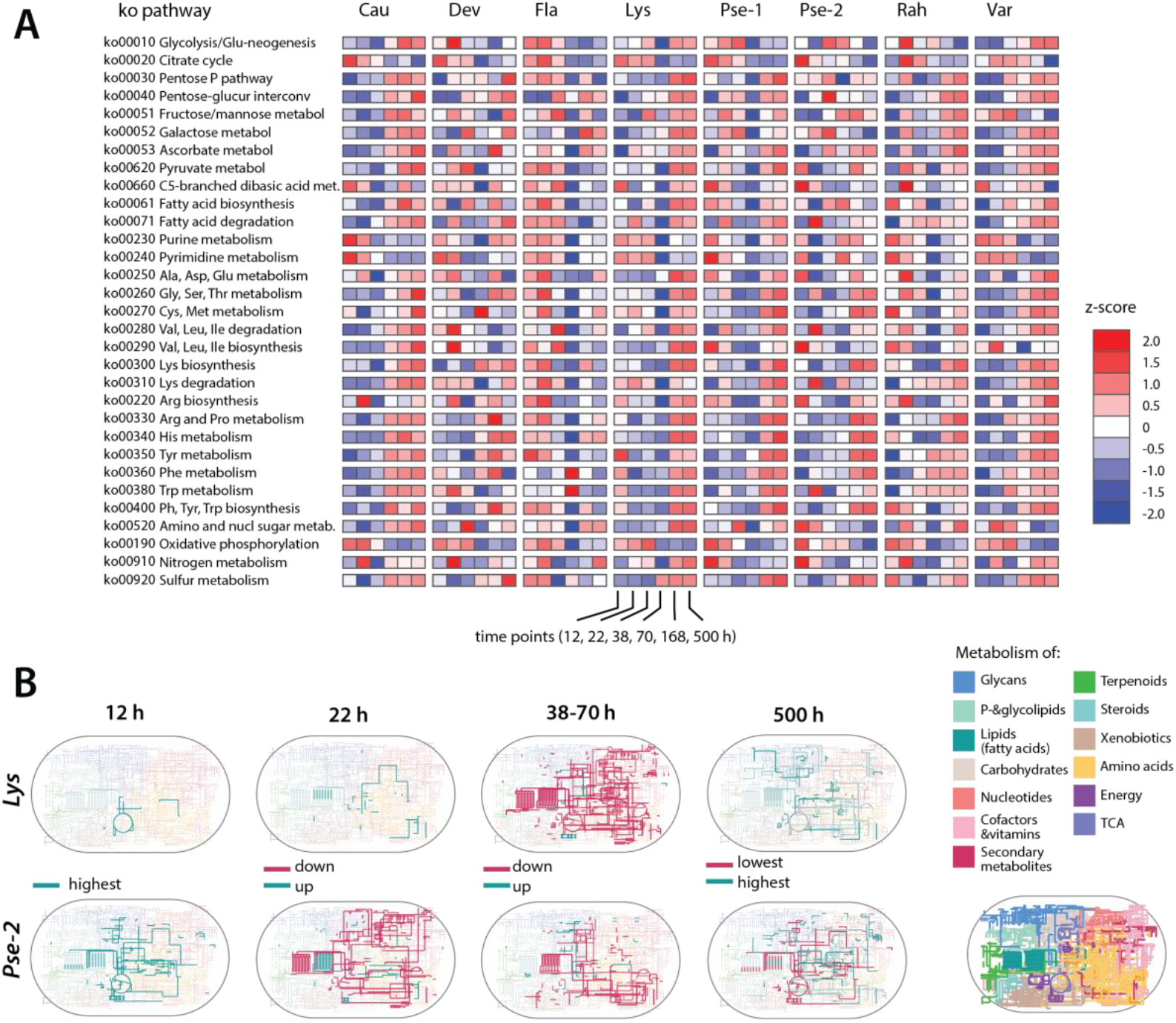
Metabolic pathway repurposing during community growth in soil. **A)** Mean expression changes in genes grouped by KEGG-orthology (ko) terms over time for the eight SynCom-21 members with reliable metatranscriptomic data (n = 4 biological replicates). Changes reported as z-score according to color legend. **B)** Changes in KEGG-pathway expression over time for *Pseudomonas* sp. strain 2 and *Lysobacter* (others, see fig. S7). Expression changes defined from categorization of time expression profiles of strain-specific genes annotated to ko-terms (as in fig. S4C & D), and grouped ko-terms per category mapped on the global KO-metabolic pathways as indicated on the right panel (colored regions correspond to metabolic pathways as per the legend).

Groups of ko-terms within the same expression category were then plotted on a global KEGG map to illustrate corresponding temporal changes in metabolic pathway expression (fig. S7). When focusing on two major community members and within the limits of resolution of the metatranscriptomics data, we can see how *Pseudomonas* sp. strain 2 becomes metabolically active faster than *Lysobacter*, with all biosynthetic pathways and genes for oxidative respiration activated at the earliest timepoint (Fig. 7B, T1 = 12 h). At the second time point (T2, 22 h), *Pseudomonas* sp. strain 2 shows growth retardation (fig. S2F) alongside decreasing KEGG pathway expression, except for fatty acid degradation and branched-chain amino acid metabolism that become induced (Fig. 7B). In contrast, all *Lysobacter* pathways are now fully active and detectable in metatranscriptomics data. A drastic change occurs in the next time points (38 and 70 h), with almost all *Lysobacter* ko-pathways showing reduced expression, including cofactors, vitamins, amino acids, sugars, and fatty acids, except for oxidative respiration (Fig. 7B, arrow; fig. S8). This change follows the peaked expression of secretion systems (Fig. 6D &E) and the biomass decrease of the assumed prey, and thus strongly suggests *Lysobacter* to uptake such compounds rather than spending more energy in renewed biosynthesis. In parallel, *Pseudomonas* sp. strain 2 has also resumed growth (Fig. 6A), but still shows decreased expression of KEGG pathways for amino acid synthesis and sugars, suggesting that at this point it is also consuming rather than biosynthesizing (Fig. 6D, T3/T4; fig. S8).

Similar individual narratives can be presented for the other main SynCom21 members (fig. S7 & S8), in some cases explaining the sharp decline in predicted growth probability at early time points (e.g., *Caulobacter* and *Devosia*’s decrease in KEGG pathway expression at 22, 38 and 70 h and decline in Fig. 6A). Certain specific KEGG-pathway expression increases point to changed substrate availability, such as the induced sugar metabolic pathways in *Devosia* at 38 h (fig. S7, arrow). *Flavobacterium’*s later population growth and decline are consistent with its KEGG-pathway expression profiles, most of which peak at 38 h but then completely collapse at 70 h (Fig. 7A, fig. S8). In summary, the metatranscriptomic data is consistent with the idea of significant substrate release from lysed cells and cross-feeding on byproducts from fast-growing populations in the community. The results also suggest that a wide spectrum of individual strategies exists, such as tuning transporters to scarce or rather abundant substrate groups, and repurposing metabolic pathways correspondingly (e.g., to avoid biosynthesis if building blocks can be acquired by import).

### Predictions of reduced membership and habitat effects on community development

Finally, we tested the model with the SMINT- and PARINT-parameter values on three different SynCom growth scenarios to understand its predictive value and/or specific limitations. In the first, we soil-inoculated reduced SynComs lacking 5-6 (rare and/or abundant) members (e.g., without *Rahnella* or Lysobacter, table S1, fig. S8). Overall, member removal caused a decrease in the fit of the simulated community trajectories compared to the full 21-member communities (Table 1), and a number of deviations in abundant populations after day 7 (fig. S8A), although overall community biomass predictions were similar to the experimental observations and better in predictions that included species interactions (fig. S8B). Subcommunities containing *Lysobacter* produced poorer overall fits and showed more variability between replicate simulations (fig. S8C), indicating a potential importance of *Rahnella* (missing in both) and *Pseudomonas* sp. strain 1 (missing in subcommunity 2) in stabilizing population development of e.g., *Variovorax, Microbacterium* or *Chitinophaga* (fig. S8C). In contrast, the subcommunity lacking *Lysobacter* was more reproducibly predicted in the simulations (fig. S8D, Table 1).

In the second scenario, we tested whether the model predictions could recapitulate an experimental dataset of growing communities with the 21 members that were weekly diluted into fresh soil (fig. S9). The advantage of this experimental dataset was that the starting community sizes and species’ relative abundances were better known, because of the tenfold dilution of the attained composition after each week. However, we noticed that several members went to almost extinction in the model simulations, but remained present in the experiments. This points at a dynamic development in the dilution-transfer experiment that the current model trained parameters do not capture well, except for a few dominant community members and which may be the result of bottlenecks in soil-to-soil transmission of rarer species (fig. S9). Indeed, the model is sensitive to variations in starting inoculum composition (both absolute and relative abundances), which can produce non-intuitive differences in growth outcomes for dominant, fast-growing and lowly abundant, slow-growing populations (fig. S10).

Finally, we retrained the SAO-algorithm on an experimental dataset of the 21-member inoculum grown in liquid soil extract in mixed suspension (Fig. 1C, fig. S11). Under these growth conditions, we would expect different interactions to emerge because of the complete mixing and all cells being exposed to a mean global change in substrate and byproduct concentrations. New SAO-optimization produced goodness of community fits similar as for the soil-grown 21-member communities (Table 1), but showed different byproduct producers and their importance (e.g., *Pseudomonas* sp. strain 1 instead of 2, fig. S11A), which was reflected in the interaction matrices (fig. S11D). In summary, these model simulations predicted general trends correctly across scenarios, although – as expected – different inoculum compositions and different habitats resulted in altered interaction patterns, which are captured imperfectly with parameters optimized for the 20- and 21-member communities grown in soil.

## DISCUSSION

We can draw two major conclusions from our work. First, we show that it is possible to parametrize two types of species interactions (SMINT, substrate-mediated byproduct formation and reutilization, and PARINT, parasitic or predatory release of cell building blocks) by iteratively minimizing discrepancies between experimental community data and simulated community growth from monoculture-derived kinetic parameters (Fig. 1, community demography model). For successful parameter estimation the experimental community growth data must encompass both changes in the absolute community size (cell numbers or biomass) and relative compositional changes over time. We demonstrate that optimized interaction parameter values are not simply fitted values without biological meaning, because (i) multiple independent initiations of the SAO-algorithm lead to converging values, (ii) parametrizing SMINT-values by random sampling of individual species’ growth rates do not result in comparable community dynamics (fig. S1), (iii) the obtained interaction parameter values can be used in a predictive manner with accurate results (Fig. 6), and (iv) the parameter values vary for communities grown in different habitats and subcommunities with different compositions are more poorly predicted (fig. S9-11). This is consistent with the concept that species interactions will change depending on resource conditions, habitat and community composition.

An important outcome of optimization by the SAO-process is that a single set for each of SMINT- and PARINT-parameter matrices is sufficient to predict community growth, independent of time. The interaction parameters in the community demography model (see *Methods* **eq. 1 & 2**), notably 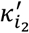 and 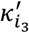, thus behave analogously as the µ_max_ and K_S_ of single population growth in relation to the (limiting) substrate condition. In analogy to µ_max_ and K_S_ changing as a function of substrate, interaction parameters change in dependence of growth conditions, and the number and constellation of neighbouring species. Our data further demonstrate that even in absence of precise knowledge on the exact resources in the soil microcosms (e.g., Fig. 2E), the SAO-algorithm can search interaction values starting from individual monoculture substrate-growth rate distributions (Fig. 2B) that can accurately reproduce and predict observed community composition changes over time. Having a generalized growth kinetic framework for microbial communities from first principles of defined parametric species interactions that link to individual growth kinetics is, to our knowledge, an important step forward.

Our second conclusion from the comparison of experimental community growth and simulations is that a staggering part (70-80 %) of community formed biomass seems to be the direct consequence of byproduct utilization through SMINT- and/or PARINT-processes, and not primary resource utilization. There are lively discussions in current literature on whether interactions among community members are on average more positive or more negative (*39, 40*), but our results indicate that this is perhaps not the right question to ask, when explaining community growth in a wider context from kinetic principles. We find that community growth in the soil microcosms is a gradual biphasic process, with a first phase dominated by primary resource utilization and a second phase characterized by assimilation and further turnover of metabolic byproducts (or, in extremo, lysed cells). Comparison of individual and community growth in soil microcosms (as in Fig. 2C) indicated that several strains profit from newly generated resources that are otherwise not available to them in the soil habitat. This byproduct contribution can be estimated by the community demography model by setting SMINT- and PARINT-parameters to zero, which indicates that there is a major positive contribution of byproduct reuse (approximately 80%) on the formed community biomass, and thus, that growing in a community for bacteria is overall profitable. We acknowledge that this behavior could be something specific for the selected soil bacteria, or for a closed system as in the microcosms, in terms of mobilizing unknown or poorly available resources that can only be achieved by a collaboration of strains. This general positive effect would have to be tested for microbiomes in other habitats. In any case, the communities of 20 or 21 isolates we worked with here are not a random assembly of laboratory strains but a representation of major soil phyla members, and, as we have shown before, their attained community compositions resemble those of typical soils (albeit of reduced diversity) (*33, 34*). For this reason, we believe that the community interaction types and magnitudes we observe and postulate have relevance for the development of natural (soil) communities. In that sense, we can conclude that species interactions have a positive effect on community growth, even though most of the strains in our defined communities are also competing for directly available carbon and nutrients.

The challenge for the community demography model as we propose it here is that there is no *a priori* knowledge of any species interaction parameter values. Although numerous methods have been proposed to measure or characterize interactions between bacterial species (*28, 41-46*), there is no consensus on the best parameter to choose, nor on the actual parametric values (like there is consensus for using the maximum specific growth rate µ_max_). We show that an SAO-algorithm that searches parameter space and minimizes the discrepancy between experimental and simulated community compositions can effectively parametrize paired-interaction matrices (Fig. 1B). Interactions are then presented as the proportion of strain biomass formation attributable to paired influence (e.g., as in Fig. 5). We are aware that the searching algorithm can produce completely unrealistic parameter values and that, given the complexity of the system and the large number of possible interactions, the results may strongly depend on the initial values. We mitigated this by using biologically meaningful initial kinetic starting points based on monoculture experiments and running multiple independently started SAO cycles. In the process of parametrizing and testing the model, we then first estimated SMINT-interaction coefficients using a 20-member community without known predatory species. These parameter values were kept unchanged and applied in a second SAO-step to explain growth of a 21-member community specifically including a *Lysobacter* predatory isolate, to derive only the PARINT-parameter values and the specific SMINT-values pertaining to its inclusion. This approach gave confident overall results for validations of unseen SynCom-20 and -21 replicates. We assess this level of confidence from four statistics applied to comparisons of experimental and simulated communities. This included comparison of the total attained community biomass (e.g., Fig. 3C, 4E), the individual member biomasses (e.g., Fig. 4B, C&D), individual population growth profiles (e.g., Fig. 4F) and complete community trajectory comparisons (Table 1, theta cosine trajectory scores). From a final set of 40 optimized parameter value matrices for byproduct and predation interactions, we then randomly picked eight to predict the community development of an independent SynCom-21 growth experiment using only the different starting composition and abundances, and the monoculture growth kinetics (Fig. 6). This resulted in very similar statistical confidence of the same comparative measures on the community behavior (Table 1).

In conjunction, paired metatranscriptomic growth state classification and expression analysis confirmed the dynamics of individual populations within the SynCom-21 (e.g., Fig. 6A), and highlighted the types of metabolic changes underlying stages in community development (Fig. 7). As the simulations also predict which community members are acting as net producer, consumer or prey at any point in time, we can quantify this contribution in terms of biomass growth or loss (Fig. 5 and Fig. 6C). In that respect it was remarkable that the simulations predicted a biphasic growth for fast-growing community members such as *Pseudomonas* sp. strain 2 and *Rahnella*, which coincided almost exactly with a transient decrease in the probability of growth predicted by mTRAC metatranscriptomic marker gene expression (fig. S2E) (*35*). Further major interactions that the model predicted between *Pseudomonas* sp. strain 2, *Lysobacter, Flavobacterium* and others (Fig. 6C) turned out to be biologically complex, with different strains having different strategies to profit from released byproducts and compounds from lysed cells (fig. S5-S8). For some strains such as *Lysobacter, Pseudomonas* sp. strain 1 and 2, and *Rahnella*, this entailed an almost complete temporary shutdown of metabolic pathway expression to take up the wealth of suddenly available building blocks (Fig. 7). Overall, the results of different independently started training cycles, comparison to randomized SMINT-values and model predictions on unseen and independent experimental data give us confidence that the obtained interaction parameter values have true biological meaning.

The community demography model is thus realistic, but, evidently, not perfect. Simulations indicated that differences in starting abundances and composition of the inoculum have a direct influence on the population succession and final biomass outcomes. This is a non-linear effect that influences abundant, fast-growing populations in a different way than rare, slow-growing members. To experimentally determine the exact starting abundances of the strains in the inoculum in the microcosms was more challenging than anticipated, because sequencing methods such as 16S rRNA gene amplicon analysis that we used to quantify relative proportions, do not take cell viability into account. In addition, it is technically difficult to quantify absolute abundances from introduced low numbers of inoculum into a soil matrix, because of extraction losses. We also saw that, although the model scaled relatively well from 20 to 21 species while keeping most SMINT-values and adding new PARINT-parameter sets, it did less well in predicting subcommunities that lacked 5 or 6 members of both dominant and rare contributors from the trained SMINT- and PARINT-sets (fig. S8). This suggests that the interaction matrices are not simply scalable by inclusion of zeros for absent members, but indeed, that the interactions themselves also change. Finally, there is also an influence of the habitat on the community developmental trajectory. This became evident from comparisons of simulations of SynCom-21 growth with soil extract in mixed liquid growth conditions and with soil extract added to soil microcosms that showed altered interaction matrices (fig. S11). This is not unexpected, but limits the extrapolation from a single parameter data set estimated for a specific community and habitat conditions.

Even though the proposed mathematical principle of biomass formation through reutilization of released waste from leaked metabolites or lysed cells is relatively simple, its complexity arises from the seemingly multitudes and varying ways that strains within communities interact and change dynamically. To improve and generalize predictive outcomes, one possibility is to generate more community growth data and to parametrize SMINT- and PARINT-interaction coefficients for specific strain compositions in the SAO-procedure within the community demography growth model (Fig. 1). This is a similar problem as measuring monoculture growth kinetic parameters on a single substrate and applying these in other conditions, but the challenge in the community demography is to obtain relevant experimental data sets with (ideally) high temporal resolution of absolute compositional changes (growth and decline). Another possibility might be to build extended genome-scale metabolic models (GEMs) which can predict resource utilization and perform dynamic flux balance analysis to predict monoculture growth rates under specific conditions (*47, 48*). Such predictions could help providing the necessary starting values to initialize our model (like in Fig. 2A &B). Newer GEM approaches are also able to simulate the overlapping metabolite release and uptake space in *in silico* communities (*28*), which could eventually provide a basis to more easily parametrize interaction coefficient matrices of the community demography model.

## METHODS

### Mathematical concept of bacterial community demography in finite systems

To describe the bacterial community demography of *N*_*S*_ bacterial species over time (i.e., their individual population growth and decline within the community) in a finite but pristine habitat (here: representative for a soil microcosm), we follow first principles of Monod growth physiology for biomass formation and waste production from resource utilization, extended by cross-feeding on reciprocal wastes as previously formulated for a two-species community (*21*). For simplicity and without other *a priori* knowledge, we assume a lumped species-specific production of waste *W*_*i*_, which by definition consists of metabolic byproducts that are not utilized at a later stage by the species itself, or else they would already be included in the biomass yield on the primary resource. SMINT-interactions in the model thus only allow a species *S*_*i*_ to consume waste *W*_*j*_ produced by a different species *S*_*j*_.

The associated system of ordinary differential equations (ODEs) that describes the change in biomass of species *i* (*dS*_*i*_); the change in waste concentrations *dW*_*i*_, and *dR*_*i*_, the change in resource concentrations, is then the sum of growth on resources and cross-feeding growth on reciprocal wastes, minus cell death (eq. 1),

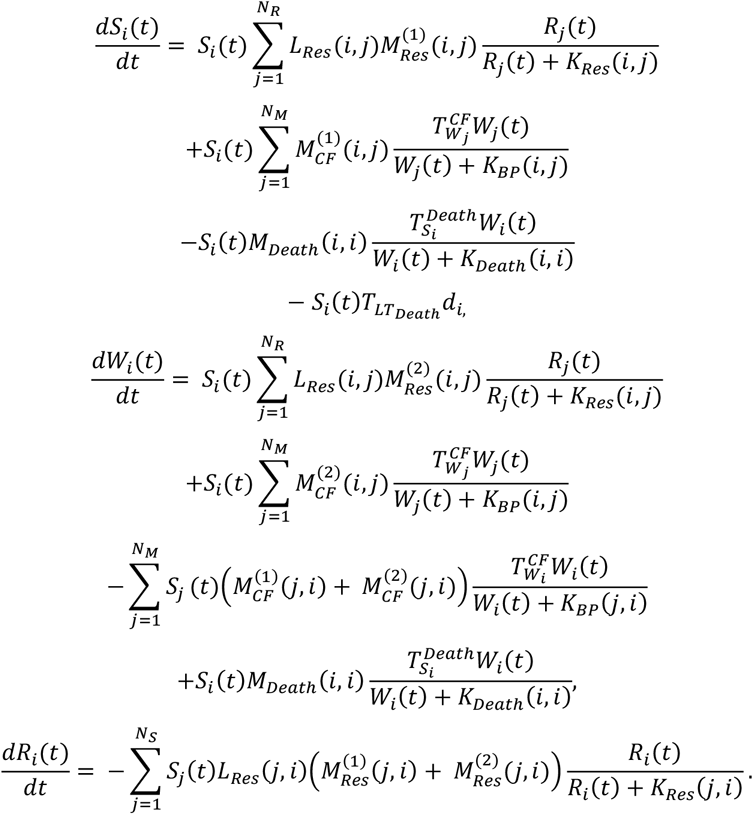

with *N*_*R*_ the number of (primary) resources in the system and *L*_*Res*_ (*i, j*) representing the lag time associated with species *i* consuming resource *j*. We introduce two resource-related matrices, 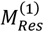 and 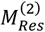, both expressed in *h*^−1^. The matrix entry 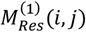 represents the effective biomass formation rate associated with the consumption of the resource *j* by species *i*, and is analogous to the Monod maximum growth rate *μ*_*max*_, or equivalently to the rate constant 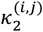. The matrix entry 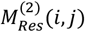 represents the effective waste production rate associated with the same process and is analogous to the rate constant 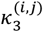. The half-saturation constant *K*_*Res*_ is defined from the underlying mass-action kinetics as 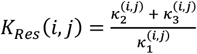, and can be interpreted as the *K*_*S*_ Monod-term of that reaction (*21*).

In similar terms, the biomass formation from byproduct *W*_*j*_ by species *i* is described by the constant 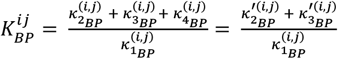 with coefficients 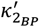 and 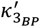 defining the interaction matrices 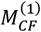 and 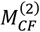 through the respective yields, and 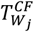 the threshold for uptake of waste. Species decay is the sum of a species-density dependent death (threshold 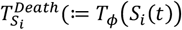) and half velocity constant *K*_*Death*_), and a constant death rate *d*_*i*_ imposed when growth substrates (*R*) become limiting below a threshold 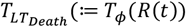. This long-term death rate (*d*_*i*_, ∀ *i* = 1, ⋯, *S*) is represented by the chemical reaction *S*_*i*_ → ∅, therefore leads to a decrease in the total biomass of the system. Thresholds have unitless values in the interval [0 1].

We assume the soil microcosm as a system with *N*_*R*_ different primary resources. Resource consumption by the *N*_S_ species is then described by the matrices 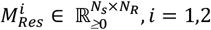 where rows correspond to the consumers (bacterial species) and columns are the resources present in the system. *N*_*M*_ denotes the number of species-specific waste products, which is equivalent to the number of species in the system since we only assume a single lumped waste per species. The species’ lag times *L*_*Res*_ for primary resource consumption are described by a matrix *N*_*S*_ × *N*_*R*_ taking value in ℝ_>0_ (units in *h*). SMINT parameters (i.e., the consumption of reciprocal waste from substrate-mediated interactions) are captured in a matrix 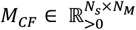 where rows are the consumer species and columns are the species-specific wastes. Because cross-feeding on waste leads to biomass formation and production of further waste, the coefficients of the cross-feeding matrix are divided into two sub-matrices. The first, 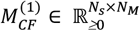 can be either positive or zero and corresponds to the rate of bacterial biomass production on reciprocal waste. The second sub-matrix, 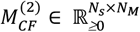 corresponds to the rate of (new) byproduct production from reciprocal waste utilization and is also positive. As explained above, species are not allowed to reuse their own waste nor new byproducts formed from waste by others, because this is implicit in their own biomass yield definition. Both matrices are related by the definition of biomass yield 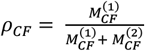. While in principle allowed that wastes could be inhibitory for growth, we specifically set 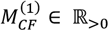 in our simulations, assuming only cross-feeding and no inhibition. The matrix *M*_*Death*_ describes the species decay (*M*(*i, i*)_*Death*_ ∈ ℝ_>0_, *M*(*i, j*) = 0 ∀*i* ≠ *j, i* = 1, ⋯, *N*_*s*_), composed of the density-dependent death and the constant turnover (see **eq. 1**).

Parasitic interactions (PARINT), here for the communities containing *Lysobacter*, are added in a matrix *M*_*Pred*_. As for the SMINT-effects, the PARINT-effect is also split in two parts 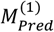 and 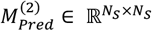, with the first explaining the rate of prey biomass being turned into predator biomass, and the second the byproducts formed from predated biomass. This implies a biomass decrease for the preys following the rate 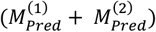 and a biomass increase for the predator following the rate 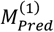. Both are related again by the predator yield ratio *α*_*pred*_ with 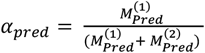. The remaining prey biomass is converted into byproduct at rate 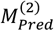 corresponding to 1 − *α*_*Pred*_. For the predation case, the complete system of ODEs in eq.1 is updated as follows (eq. 2),

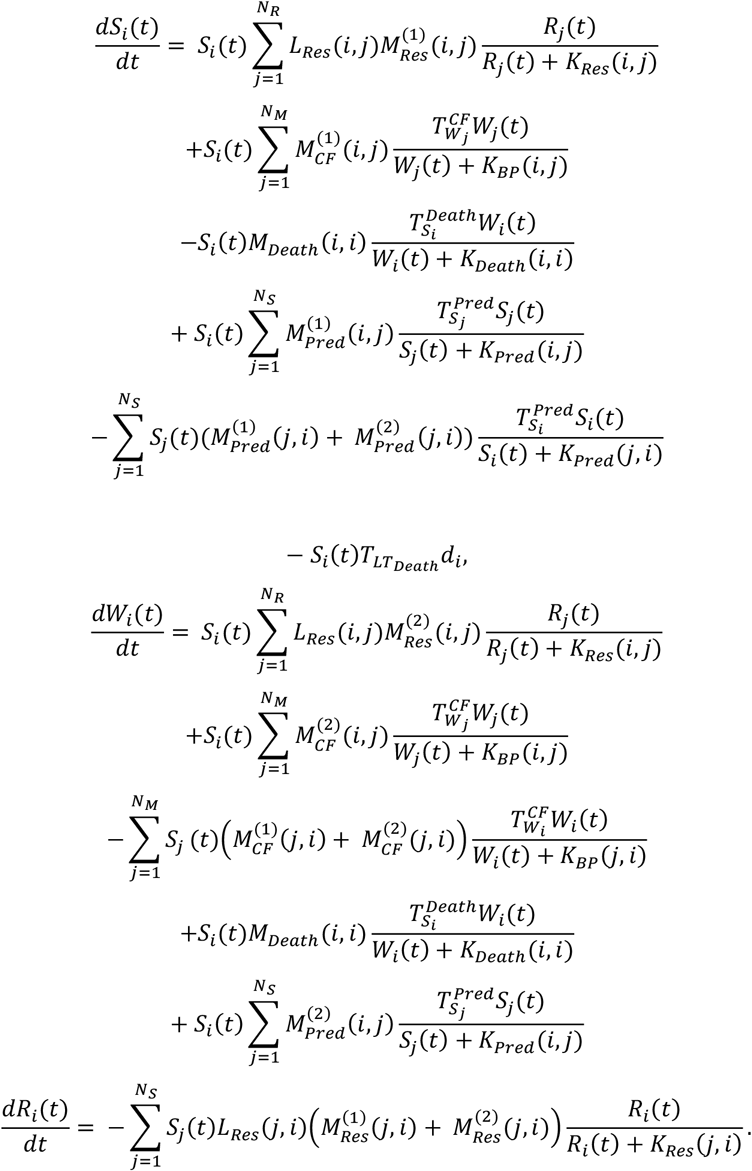

The cross-feeding threshold 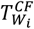 depends on byproduct concentration, and the threshold function 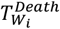 depends on the concentration of a species *S*_*i*_. To be consistent with the previous notations, we use the notation *S*_*i*_ = *X*_*i*_ within ODEs to identify the total biomass of a bacterial species *S*_*i*_, but also as a reference for the species itself. The ODE-implicit chemical reaction network implies a total (bio-)mass conservation within the system with finite boundaries.

To implement the model for *N*_*S*_ species and their SMINT-parameters, the ODEs of **eq.1** are executed on a set of matrices (Algorithm 1), which are individually and sequentially solved as outlined in Algorithm 2. In case we include PARINT-parameters, we execute the ODEs of **eq.2** and include the *M*_*pred*_-matrices. When searching only PARINT-but not SMINT-interactions, matrix values from previous SMINT-searches can be kept. The order of matrix layer fitting in the algorithm should theoretically not matter, but might affect the results depending on the number of iterations.

#### Algorithm 1 Community demography matrices

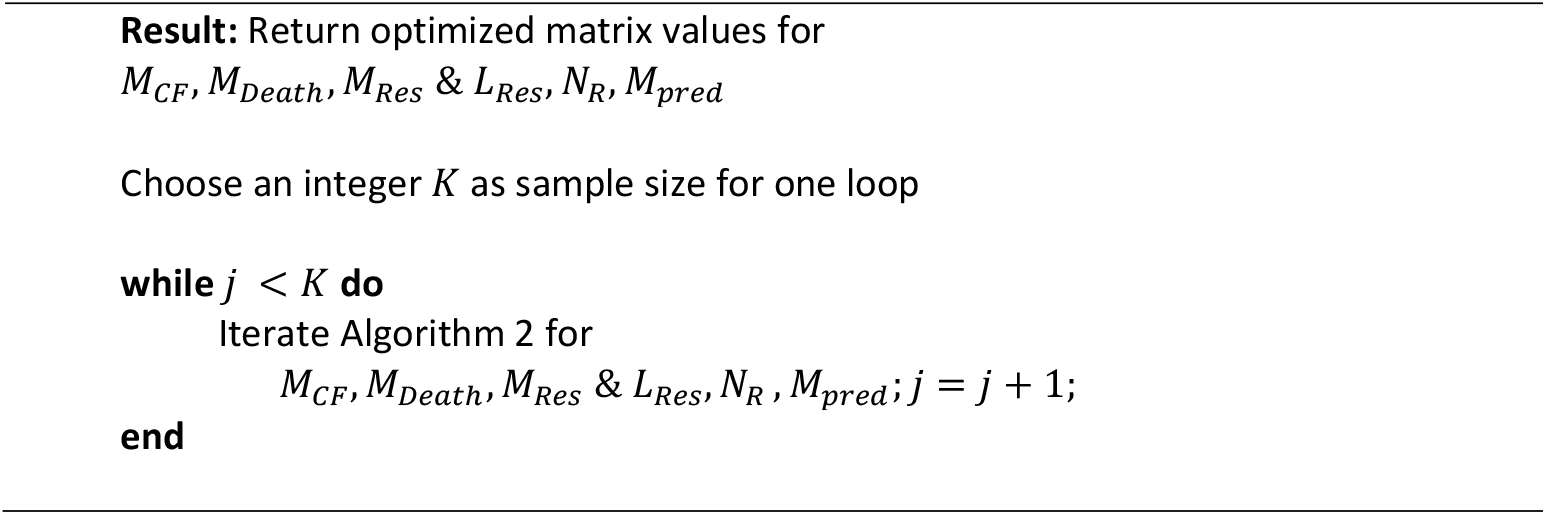

### Matrix parameter estimation from simulated-annealing optimization iteration

Depending on the chosen community composition (20 or 21 species, with or without predator; see below), we preset or optimize selected parameter values in the matrices of Algorithm 1 using a simulated annealing optimization (SAO) algorithm combined with an adaptive Gaussian random-walk proposal inspired by the robust adaptive Metropolis scheme (*49*). The algorithm is divided into four blocks. In each block, an SAO procedure is applied to sequentially fit the cross-feeding, self-inhibition, primary resource consumption, and predation interaction parameters, while keeping the other interaction parameters fixed. After the last block, the procedure returns to the first block to perform a second round of fitting. This process is repeated until a predefined number *K* of block iterations is reached. At each iteration, candidate parameter matrices are generated using a multivariate normal perturbation whose covariance is adapted to target a desired acceptance rate. The candidate parameters are then used to solve the ODE system of eq. 1 (in case of only SMINT-interactions) or eq. 2 (in case of SMINT- and PARINT-interactions), and the resulting simulated community compositions are compared to the empirical values through a weighted error-based objective function (Algorithm 2).

Candidate solutions are accepted with probability:

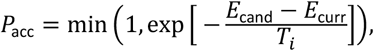

where *E*_cand_ and *E*_curr_ denote the objective function values of the candidate and current parameter sets, respectively, and *T*_*i*_ is a temperature parameter that decreases over iterations according to a predefined decay schedule. High temperatures allow wider exploration of the parameter space, while progressively lower temperatures concentrate the search around low-error solutions. Although the candidate value proposal distribution and covariance adaptation follow principles of robust adaptive Metropolis sampling, the algorithm is used here exclusively for optimization rather than Bayesian inference. Parameter sets of growth rates, resource consumption rates, and cross-feeding rates are further constrained to the interval [0 10] h^−1^ to avoid unrealistic values. Death rates are allowed to vary between 0.012 and 10 h^−1^.

Matrices are initiated as follows. The initial starting rates of the individual species in the community in *M*_*Res*_ are randomly sampled from their log-normal fitted distribution of maximum observed growth rates across a range of substrates and substrate conditions (Fig. 2B; see below). Similarly, the initial lag times in *L*_*Res*_ are randomly drawn from their log-normal fitted lag phase distributions extracted from the same monoculture growth conditions as for μ_*max*_. The initial rates in the *M*_*CF*_ cross-feeding matrices are independently sampled from the log-normal maximum observed growth rate distributions of the monocultures but multiplied by 0.8, under the assumption that byproducts are used less efficiently than the primary resources. The *M*_*Death*_ matrix is initiated with an equal death rate for each species of 0.0102 h^−1^ as suggested by literature values (*50*). As initial value for the predation rates in *M*_*Pred*_ we use 0.01 h^−1^. Based on targeted chemical analysis of aqueous-phase extracted compounds from the soil microcosms we set the number of resources in *N*_*R*_ to 12, and divide the total measured organic carbon (TOC) concentration in the soil microcosm system among the 12 compound groups in proportion to their summed peak areas in the chemical analysis to obtain their starting concentrations (Fig. 2E, table S1). For fitting scenarios we take the starting biomass (in g C DW ml^−1^) of the individual species in the community as the mean of the experimentally determined absolute cell abundances (for each of the species) at t=0, multiplied by the average cell dry weight calculated according to Ref. (*51*).

#### Algorithm 2 Matrix solver

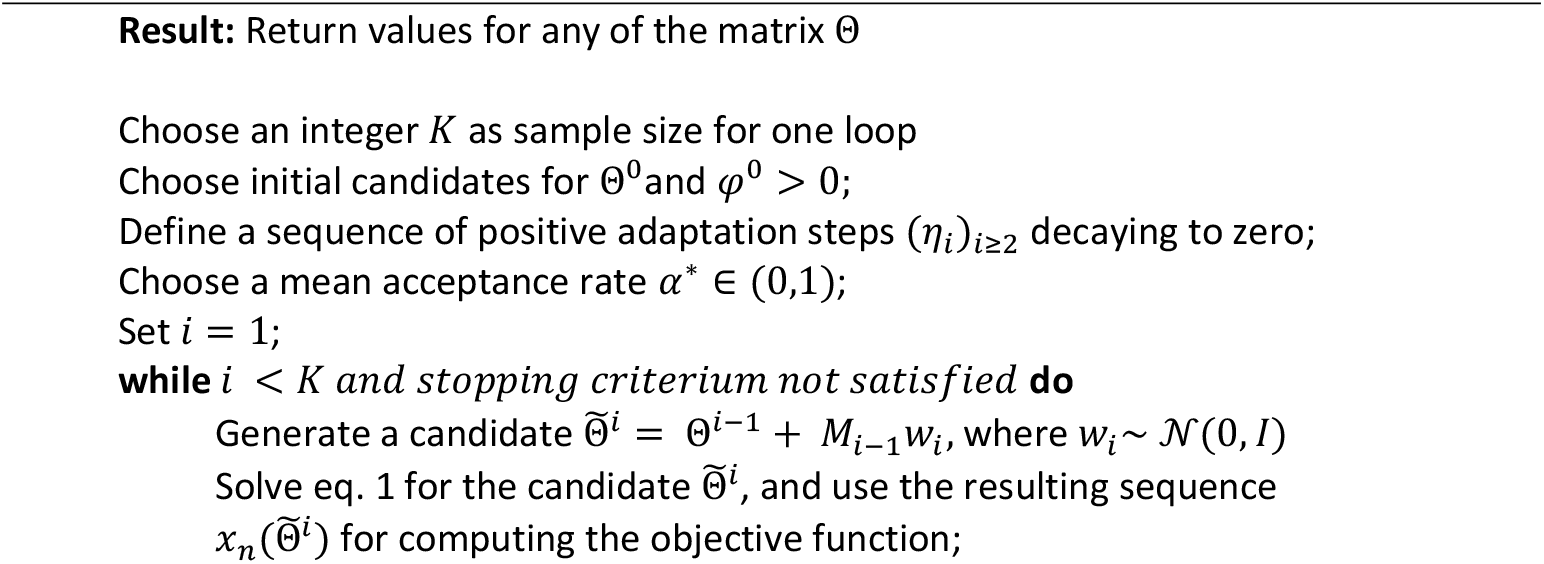
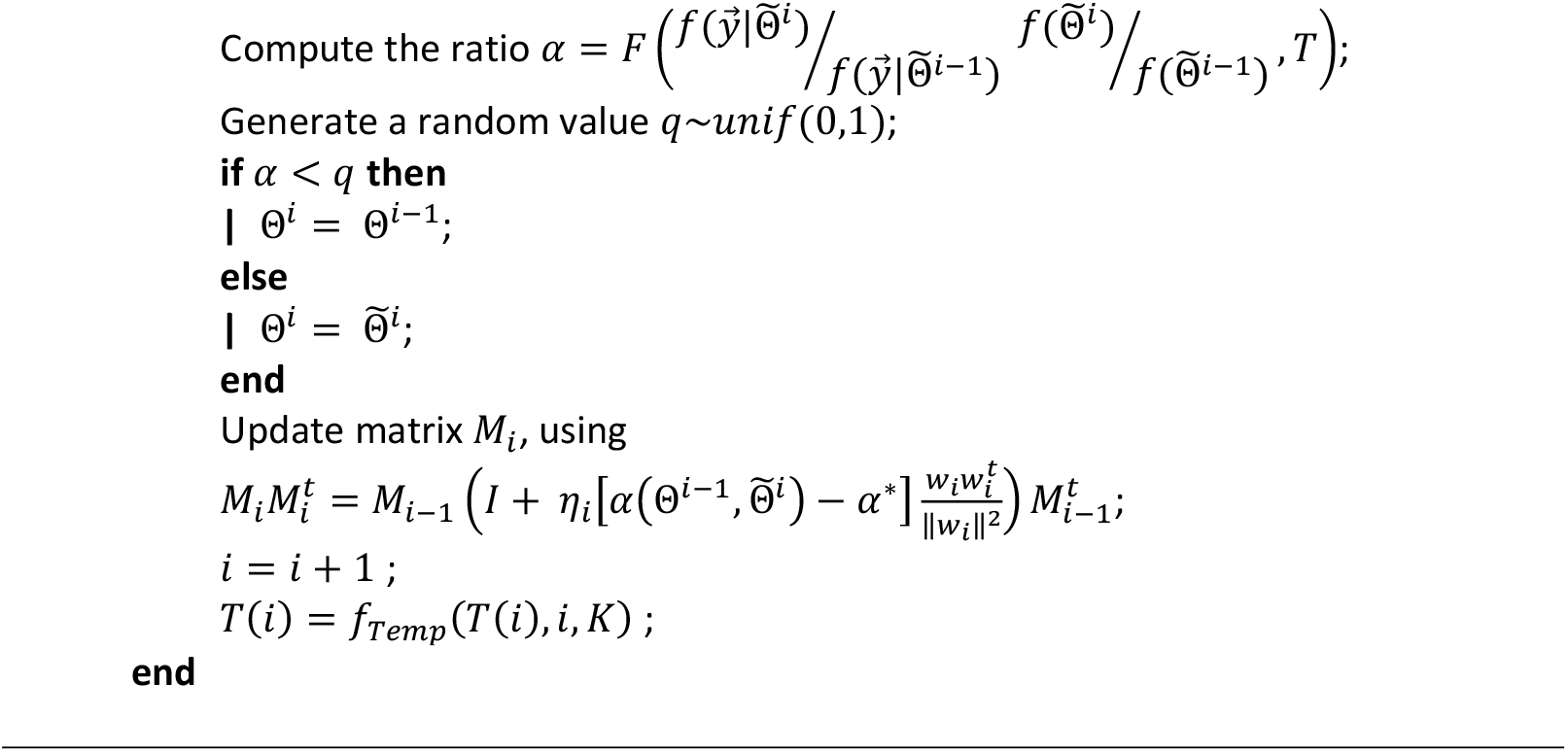

### Community demography simulation training and prediction scenarios

To test and parameterize the community demographic model using experimental data, we constructed bottom-up synthetic communities of either 20 (SynCom-20) or 21 bacterial species (SynCom-21, same members as SynCom-20 but additionally, the facultative bacterial predator *Lysobacter*, table S2) cultured in soil microcosms and sampled over a period of three weeks (see below and Table 1 for the specifics of each experiment). Each experiment produced the relative abundances over time of the individual populations forming the community (from 16S rRNA gene amplicon sequence data, with the soil-purified bacterial cell fraction as basis for DNA isolation; see below) and the total cell counts of the isolated cell fraction. By multiplying the relative abundances with the total cell counts we derived absolute population abundances over time (assuming that there were no species-specific DNA isolation biases). Each experiment was conducted in 4-5 biological replicates.

To estimate SMINT-parameter values and test their effects, we used data from microcosm-grown communities of 20 soil isolates (SynCom-20), without *Lysobacter*. Two independently repeated microcosm series, each with n=4 biological replicates were combined and split into two parts: one part used for the SAO-algorithm (62.5% of the data, containing all replicates from the first experiment and one replicate from the second), and the second part used for validation (37.5% of the data). In this case, the model matrices comprised *N* = 20 species and 12 resource groups. The initial matrix parameterization followed the procedure described above, followed by the Algorithm 2-procedure to find the best cross-feeding matrix parameter sets that minimize the discrepancy between simulated absolute biomass values of each community member and the corresponding experimental measurements at each time point. This fitting procedure was repeated independently 10 times, yielding 10 independent parameter sets, which were compared to assess the uniqueness and consistency of the optimization process when starting from the same initial conditions.

As a control for the importance of SMINT-parameter optimization, we pick one of the 10 fitted *M*_*Res*_ and populate the *M*_*CF*_ matrices per consumer-species with randomly sampled values from their log-normal growth rate distributions multiplied by 0.8. This was repeated six times (fig. S1).

To parametrize PARINT-effects (i.e., including the additional *M*_Pred_ matrix as explained above in eq. 2), we used soil-microcosm grown communities of 21 strains, now including *Lysobacter* (*n* = 4 biological replicates; labeled SynCom-21 in Table 1). Importantly, this model simulation uses one of the optimized sets of the *M*_CF_ cross-feeding matrices values of the 20 species from the SynCom-20 in **eq. 1**. The Algorithm 2 procedure then only optimizes the resource/byproduct consumption and death parameters for *Lysobacter* itself, and the predation matrices for the inclusion of *Lysobacter* in SynCom-21. We bias the potential preys for *Lysobacter* in the matrices based on a comparison of the species biomass between the SynCom-20 and SynCom-21, resulting in only *Caulobacter, Flavobacterium, Microbacterium*, and *Variovorax*. Additionally, we assume a yield of *Lysobacter* biomass formation from predation of 20%. With this simulation strategy the number of parameters to be estimated was drastically reduced, limiting risks of overfitting. This PARINT SAO-procedure was repeated 40 times, yielding 40 combined matrix sets of SMINT- and PARINT-parameter values.

SMINT- and PARINT-matrix parametrization was repeated independently for SynCom-21 community data growing in mixed-suspended liquid culture on soil extract medium (fig. S11).

Finally, we use 10 randomly drawn from the 40 optimized SMINT- and PARINT-parameter value sets from soil-microcosm data to predict the outcomes of further defined communities grown in soil microcosms (Fig. 1C). First, we repeated an independent SynCom-21 experiment (labeled SynCom-21 *MT* for the Meta-Transcriptomics inclusion, Table 1) that was started from a freshly prepared mixed inoculum. Because the first time point of the SynCom-21 *MT* experiment was only taken after 12 h, we used starting abundances for each of the community members that were retroprojected to t=0 from the experimental data at t = 12 assuming exponential growth. Secondly, we produced three strain drop-out variants of the 21-member defined community without rare or abundant members (fig. S8), which were regrown in sterile soil microcosms as the other communities. Third, we use previous community data of weekly-transferred and one-tenth diluted soil microcosms inoculated with the 21-member defined community (*34*) to *de novo* predictions of species growth.

### Soil microcosm community growth experiments

SynCom-20 and SynCom-21 inocula (table S1) were cultured in sterile soil microcosms consisting of 90 g dried, double-autoclaved river bank matrix (mostly sand and silt) in 500 ml capped Schott glass bottles, complemented with 10 ml of a sterile water-extract of forest top soil (soil extract) to provide additional carbon and nitrogen, as previously described (*34*). The measured total organic carbon content of soil extract amounts to 0.738 mg C per ml, and the microcosms were kept at 10% gravimetric water content. Microcosms were inoculated with strain monocultures, or with the mixture of 20 or 21 species and subsets thereof, in approximately equivalent cell counts, each individually precultured on R2A agar plates (*52*) for 5 days at 18–20 °C and harvested using a stainless-steel spreader bar and 5 ml pyrophosphate solution (0.2% *w/v* tetrasodium-pyrophosphate decahydrate solution at pH 7.5). Harvested cells were suspended gently by pipetting and then centrifuged at 4000 × *g* for 7 minutes. After removing the supernatant, the cells were resuspended in 5 ml pyrophosphate solution. Cell densities were SYBR-Green-stained and quantified by flow cytometry as previously described (*34*) and diluted, if needed, to a density of 10^9^ cells ml^−1^. Aliquots of 1 ml of each strain were then pooled together, mixed by slow vortexing and diluted with sterile soil extract to produce an inoculum of 10^7^ cells ml^−1^. Four replicate microcosms (90 g matrix) were inoculated with 10 ml of the inoculum to achieve a starting density of 10^6^ cells g^−1^ soil. DNA was isolated from the inoculum (see below) to characterize the starting relative abundances of each strain. Inoculated glass bottles were capped and placed on their sides on a Stuart SRT9D horizontal roller mixer (Cole-Palmer, UK) during 2 h at room temperature to distribute the moist soil-inoculated material homogenously. Microcosms were then incubated upright in the dark at 18-20 °C for a duration of up to three weeks.

Microcosms were sampled at defined time intervals (typically, after 1 day, 3, 7, 9 or 10, 15 and 21 or 22 days) to quantify the community size and its composition. Before sampling, the bottles were placed on a horizontal roller mixer for 30 minutes, after which 10 g soil was removed with a sterile spatula (weight recorded) and transferred to a 15 ml sterile Falcon tube (and the remainder microcosm was incubated further). Microbial cells were detached from the matrix by vortexing the soil samples with 10 ml 0.2% sodium pyrophosphate solution on a Vortex-Genie 2 (Scientific Industries, USA) at full speed for 2 min. Tubes were then placed vertically for 3–5 minutes to sediment particles, and 5 ml of the clear liquid phase above the sedimented material was aspirated and transferred into a fresh 15 ml Falcon tube placed on ice. A 600–μl aliquot of this cell suspension was further passed through a 40–μm cell strainer (Corning, USA) into a 2 ml Eppendorf tube and placed on ice for quantifying total cell counts by flow cytometry. The remaining cell suspension was centrifuged at 4000 rpm for 7 minutes at 21 °C in a swing-out rotor 5810R centrifuge (Eppendorf, USA). The supernatant was removed and the pellet was resuspended in 2 ml pyrophosphate solution, which was transferred to fresh 2 ml Eppendorf tubes. This suspension was centrifuged at 7000 × *g* for 7 minutes and the supernatant removed to leave a dry cell pellet. These cell pellets were stored at –80 °C until total DNA extraction.

### 16S rRNA gene amplicon sequencing and analysis

Total DNA was extracted and purified from thawed cell pellets with a DNeasy PowerSoil Pro Kit (Qiagen, Germany) according to the provided protocol. Quantities of 10 ng purified DNA were used to amplify the V3–V4 region of the 16S rRNA gene in the polymerase chain reaction, which were subsequently indexed using the Nextera XT primer 1 Set A/B and corresponding protocol as per instructions by Illumina, described in detail in (*53*). The indexing strategy allowed for a final multiplexing of 192 samples per library, which were spiked with 30% PhiX control v3 (Illumina Inc., USA) prior to 300 cycle paired-end sequencing on MiSeq v3 (Illumina Inc., USA) or AVITI Sequencing System (Element Biosciences, San Diego, CA, USA) using a Cloudbreak Freestyle™ Medium Output Kit at the Lausanne Genomic Technologies Facility.

The quality of the sequence reads was checked with FastQC 0.11.9 (*54*) and trimmed using Trimmomatic 0.39 (*55*) with a sliding window of 5:28. Paired reads were then merged using Flash 1.2.11 (*56*) with a 20 bp minimum and 170 bp maximum overlap with no allowed mismatches. To match reads to the members of the SynCom-20 or SynCom-21, we collected and aligned all sequence variants from the genome of each member (table S2) using ContEst16S (https://www.ezbiocloud.net/tools/contest16s) and Kalign3 (*57*). Species-specific regions (99–100 bp) were chosen using Jalview 2.11.2.5 and used as a query for *grep*, the Bash pattern matching tool, to count the number of each sequence per sample. Absolute abundances of each query sequence matching to the target species were summed and then divided by the number of rRNA operons in each species. This number was normalised by the total number of *grep* matches per sample to calculate the relative abundance of each community member. Absolute SynCom member abundances in the microcosms were then obtained by multiplying the relative abundances from the 16S rRNA gene-derived counts with the total cell counts from flow cytometry.

### RNA and DNA isolation for metatranscriptomic and metagenomic sequencing

For transcriptomics and microbial community analysis, microbial cells were first extracted from the soil matrix by combining 10 g of microcosm with 20 ml of ice-cold 4 g/l sodium pyrophosphate decahydrate solution in 50 ml polypropylene conical tubes (Greiner bio-one) and 2 min vortexing at maximum power settings on Vortex-Genie® 2 mixer (Scientific Industries). Cells from the supernatant were collected by 5 min centrifugation at 3220 *g* in a swing out rotor (Eppendorf A-4-62) at 0°C, flash-frozen in liquid nitrogen and stored at −80°C. Total DNA and RNA were extracted using the RNeasy PowerSoil Pro Kit combined with RNeasy PowerSoil DNA Elution Kit (Qiagen, Hilden, Germany) according to the manufacturer’s instructions.

Following extraction, RNA was treated with TURBO DNAse (Invitrogen, Thermo Fisher Scientific, Waltham, MA, USA) to remove residual genomic DNA and concentrated using an RNA Clean & Concentrator-25 (Zymo Research, Irvine, CA, USA). Metatranscriptomic libraries were prepared from 100-250 ng total RNA using the Zymo-Seq RiboFree Total RNA Library Kit (Zymo Research, Irvine, CA, USA), which includes an integrated rRNA depletion step, following the manufacturer’s protocol. Libraries were indexed using the Nextera XT v2 Index Kit A (Illumina, San Diego, CA, USA). Libraries were purified using magnetic beads and pooled together, quantified with Qubit dsDNA BR Assay Kit and QuBit 2 Fluorometer (Invitrogen, Thermo Fisher Scientific, Waltham, MA, USA), and fragment size distributions were assessed by capillary electrophoresis on a 5300 Fragment Analyzer System (Agilent Technologies, Santa Clara, CA, USA) prior to sequencing. 150-bp paired-end sequencing was performed on an AVITI Sequencing System using a Cloudbreak Freestyle™ High Output Kit (Element Biosciences, San Diego, CA, USA) at the Lausanne Genomic Technologies Facility.

### Metatgenomic and metaranscriptomics data processing

The resulting raw reads from metagenomic and metatranscriptomic sequencing were cleaned by removing adaptor sequences, low-quality-end trimming and removal of low-quality reads using BBTools v 38.18 (*58*). Available from: https://sourceforge.net/projects/bbmap/. The exact commands used for quality control can be found on the Methods in Microbiomics webpage (Sunagawa, S. Data Preprocessing - Methods in Microbiomics 0.0.1 documentation. Available from: https://methods-in-microbiomics.readthedocs.io/en/latest/preprocessing/preprocessing.html [doi: https://doi.org/10.5281/zenodo.15019381]). For metatranscriptomic analysis, reads were mapped to the concatenated soil SynCom-21 genomes with bowtie2 v. 2.5.1 (*59*) and quantified using featureCounts v. 2.0.1 (*60*). Statistical analysis and data normalisation were performed independently for each strain as described in Ref. (*61*), a Python re-implementation of the DESeq2 framework (*62*). DE-genes for *Lysobacter* were further identified on log_2_-TPM-counts for which at least 3 replicate data values were available using MATTEST and MAFDR multiple test corrections implemented in MATLAB (version R2023b). Probabilities of growth were inferred from expression of sets of conserved household genes using mTRAC with default parameters (*35*). The quality-controlled metagenomic reads were used for in situ growth rate prediction of each of the members of the SynCom-21 *MT* community with CoPTR, under default parameters (*36*).

KEGG Orthology terms were assigned to predicted proteins using eggNOG-mapper v2.1.12 with default parameters against the eggNOG database (version 6.0) (*63*). Transporter annotations were generated using gapseq with find –p transporter mode (*64*). Expression changes over time and per species in the SynCom-21 *MT* experiment were further analyzed by KEGG orthology pathway terms (i.e., ko-terms) (*65*). Expression of genes annotated to ko-terms or transporters were categorized by their time profile based on Pearson similarity scores to a select ‘type’ example (fig. S4 and Fig. 4E). For direct representation, ko-pathway expression was plotted as the log_2_-TPM values per genome and time (e.g., Fig. 6D). Expression differences over all common ko-pathway annotated genes were summarized in principal component analysis (as in fig. S6). Ko-pathways allocated to the same time profile category of fig. S4C were then mapped per time-point and genome on the global KEGG metabolic maps by iPATH3 (*66*) for representation, as explained in fig. S7 and summarized for a select pair of dominant strains in Fig. 7B.

### Quantifying total cell counts by flow cytometry

Total cell counts in the recovered soil cell suspensions were quantified by flow cytometry of SybrGreen-I stained suspensions according to the procedure outlined by the supplier (Thermofisher), counting gated events above a FITC-value of 1000, as described previously (*53*). Cell counts were then converted to cells per ml of soil water phase, using a gravimetric water content of 10 ml water per 100 g total material. Cell counts were converted to average dry weight biomass (table S1) using the procedure outlined in Ref. (*51*).

### Individual species growth rates and substrate preferences

Different complementary methods were used to measure maximum specific growth rates of the individual SynCom members in pure liquid culture: (i) succinate as sole carbon substrate in minimal medium, (ii) growth on a complex substrate composed of peptone, tryptone, yeast extract and glucose (PTYG), (iii) growth on soil extract medium and (iv) substrate utilization in BIOLOG plates. Individual maximum population densities in soil measured after 1 week of growth served to calculate an average biomass yield (*n* = 4 replicates).

Growth kinetic measurements were started as follows. Individual SynCom members were recovered from their –80 °C glycerol stock, plated for five days at 18-21 °C on R2A agar, after which a single isolated colony was resuspended in 200 µl 21C minimal medium (*67*) with 5 mM succinate in a single well of a 96-well plate. The increase in culture turbidity at 600 nm was measured using a BioTek Synergy H1 plate reader for 48 h, with four replicates for each strain. In case of PTYG, the procedure was repeated with medium containing 0.6 MgSO_4_·7H_2_O, 0.07 g CaCl_2_·2H_2_O, 0.5 g peptone, 0.5 g tryptone, 1 g yeast extract and 1 g glucose per L. For growth on soil extract medium, strains were inoculated at similar densities as before, but in this case, we measured population growth from increase in flow cytometry total counts. For BIOLOG experiments, each strain was tested in triplicate on 30 different carbon substrates, and the respiration values from conversion of the resazurin salt were measured using a BioTek plate reader at 590 nm. Growth curves were fitted with a Monod substrate-to-biomass function to extract maximum population size, *K*_*S*_, lag time and μ_*max*_, as described previously (*21*).

### Liquid chromatography mass-spectrometry metabolite analysis

In order to get an approximation of the substrate utilization in the soil microcosms by the growing SynCom-21, we sampled (5 g) sterile control soil microcosms with soil extract added, and three replicates of inoculated SynCom-21 microcosms after day 1, 3 and 10. Soil aliquots (5 g) were resuspended in 25 ml pyrophosphate solution, vortexed at 700 rpm for 2 min, settled for 1 min, after which 0.5 ml of supernatant was transferred to a clean Eppendorf vial. This was centrifuged for 2 min at maximum speed (15,000 rpm in a Biofuge), the supernatant was filtered over a 0.2 µm syringe filter and flash-frozen in liquid nitrogen. Samples were stored at –80 °C until analysis by Hydrophilic Interaction Liquid Chromatography coupled to tandem mass spectrometry (HILIC – MS/MS) at the University of Lausanne Metabolomics Facility. As this is targeted MS, we represent compounds by their peak areas. Compounds are grouped by chemical category (table S2) to estimate 12 substrate classes in the soil (and soil extract liquid) that build the *N*_*Res*_ matrix mentioned above. We calculate the relative proportions of the total by their peak area, sum those per chemical category, and use that proportion to derive an equivalent compound availability (in mg C l^−1^) by multiplying the proportion with the TOC concentration of soil extract.

### Statistical analysis

To assess the quality of the total biomass predictions between different models and simulations, we use Cliff’s delta, a nonparametric effect-size measure. Cliff’s delta quantifies the degree of difference between two groups without assuming any underlying distribution and does not require a minimum sample size. The statistic ranges from −1 to 1, where 0 indicates complete overlap between the two groups, and values closer to ±1 indicating that one group consistently yields larger values than the other. A statistical significance can be attributed to Cliff’s delta by permutation testing (n = 100,000 permutations). Group labels are randomly permuted after pooling the two observed groups, Cliff’s delta is recomputed for each permutation, and the *P* value is reported as the proportion of permuted statistics that are at least as extreme as the observed value.

To assess similarities in community composition at the last time point, we performed a PERMANOVA test on the Bray–Curtis distance matrix. Assuming we have *N* replicates of a community with *S* species, the Bray–Curtis distance matrix is defined as,

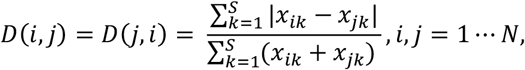

where *x*_*ik*_ denotes the abundance of species *k* in replicate *i*. Here, *D*(*i, j*) = 0 corresponds to identical community composition, whereas *D*(*i, j*) = 1 indicates completely different community compositions. The PERMANOVA permutation test evaluates the null hypothesis that the centroids and dispersion of the communities are equivalent between the considered groups.

To assess the similarity between observed and simulated growth of individual populations, we use the Pearson correlation coefficient and its associated *P* value. The cosine similarity index was computed to compare community trajectories at each time point and averaged over all time points. Values close to 1 indicate similar dynamics, values close to 0 indicate different dynamics, while negative values correspond to opposite dynamics. We complement these values with a permutation test to evaluate the significance of the obtained similarity score.

## Data availability

The community demography model (MATLAB version 2023b) is available from a single Zenodo link (Version v1.0.0; 10.5281/zenodo.20428510) and from GitHub (https://github.com/IsalineLucille22/Biphasic-bacterial-community-assembly-predicted-from-generalized-first-principles-of-monoculture-gro).

The raw sequencing data generated in this study have been deposited in the European Nucleotide Archive (ENA) under accession number PRJEB112616.

## Acknowledgments

The authors thank Alan Pacheco for helpful critical comments on the manucript, and all members of the NCCR Microbiomes for stimulating discussions during the various stages of this work. We thank Davin Wijaja for his help in genome assemblies of the SynCom-21 isolates.

Parts of the abstract were improved for clarity by use of large-language models and verified by human authors.

## Author contributions

I.G., C.M., S.C., M.S., A.S., S.S. and J.M. conceived the studies and designed experiments. S.C., C.B., V.S. conducted soil microcosm experiments. M.S., A.S., H.R. and J.M. analyzed metagenomic and metatranscriptomic data. I.G., A.V. and C.M. conceptualized and coded the community demography model. I.G. and J.M. conducted model simulations. I.G., C.M., A.S. and J.M. wrote the draft manuscript. All authors provided input, verified, corrected and/or consented the final manuscript.

## Supplementary information

table S1: Characteristics of the 21 soil isolates.

table S2: HILIC analysis of soil water extractable compounds.

Supplementary FIG 1. Simulation outcomes of soil-growth of the 20-member defined community with random SMINT-values.

Supplementary FIG 2. Community demography in the 21-member SynCom used for meta-transcriptomic analysis

Supplementary FIG 3. Prediction of population growth in the SynCom-21 based on CoPTR calculations.

Supplementary FIG 4. Classification of transporter and KEGG-orthology pathway expression types in the 21-membered SynCom used for metatranscriptomic analysis.

Supplementary FIG 5. Predicted substrate detail for any of the Type 1 (Peak at timepoint 2 or 3) or Type 2 (troughs) transporter expression profiles

Supplementary FIG 6. Principal component analysis comparison of ko-pathway annotated expression changes among the eight abundant species in the SynCom-21 metatranscriptomics experiment.

Supplementary FIG 7. Temporal changes in KEGG orthology metabolic pathways in the growing SynCom-21 metatranscriptomics experiment

Supplementary FIG 8. Predictions from the SynCom-21 SMINT/PARINT-model for growth of subcommunities with reduced composition.

Supplementary FIG 9. Predicted (A) and observed (B) community compositions of a weekly soil-diluted SynCom-21.

Supplementary FIG 10. Simulated effect of differences in inoculum composition on community development.

Supplementary FIG 11. Changing species interactions during SynCom-21 growth in liquid suspension.

## REFERENCES

1. J. Monod, The Growth of Bacterial Cultures. Annual Review of Microbiology 3, 371 (1949).DOI 10.1146/annurev.mi.03.100149.002103

2. D. K. Button, Kinetics of nutrient-limited transport and microbial growth. Microbiol Rev 49, 270 (1985).10.1128/mr.49.3.270-297.1985

3. S. A. Kooijman, E. B. Muller, A. H. Stouthamer, Microbial growth dynamics on the basis of individual budgets. Antonie Van Leeuwenhoek 60, 159 (1991).10.1007/BF00430363

4. S. J. Pirt, The maintenance energy of bacteria in growing cultures. Proc R Soc Lond B Biol Sci 163, 224 (1965).10.1098/rspb.1965.0069

5. L. R. Thompson, J. G. Sanders, D. McDonald, A. Amir, J. Ladau, K. J. Locey, R. J. Prill, A. Tripathi, S. M. Gibbons, G. Ackermann, J. A. Navas-Molina, S. Janssen, E. Kopylova, Y. Vazquez-Baeza, A. Gonzalez, J. T. Morton, S. Mirarab, Z. Zech Xu, L. Jiang, M. F. Haroon, J. Kanbar, Q. Zhu, S. Jin Song, T. Kosciolek, N. A. Bokulich, J. Lefler, C. J. Brislawn, G. Humphrey, S. M. Owens, J. Hampton-Marcell, D. Berg-Lyons, V. McKenzie, N. Fierer, J. A. Fuhrman, A. Clauset, R. L. Stevens, A. Shade, K. S. Pollard, K. D. Goodwin, J. K. Jansson, J. A. Gilbert, R. Knight C. Earth Microbiome Project, A communal catalogue reveals Earth’s multiscale microbial diversity. Nature 551, 457 (2017).10.1038/nature24621

6. J. F. Matias Rodrigues, J. Tackmann, L. Malfertheiner, D. Patsch, E. Perez-Molphe-Montoya, N. Napflin, D. Gaio, G. Rot, M. Danaila, M. E. Peluso, M. Dmitrijeva, T. S. B. Schmidt, C. von Mering, The MicrobeAtlas database: Global trends and insights into Earth’s microbial ecosystems. Cell, (2026).10.1016/j.cell.2026.01.021

7. R. MacArthur, R. Levins, Competition, Habitat Selection, and Character Displacement in a Patchy Environment. Proc Natl Acad Sci U S A 51, 1207 (1964).10.1073/pnas.51.6.1207

8. M. Dal Bello, H. Lee, A. Goyal, J. Gore, Resource-diversity relationships in bacterial communities reflect the network structure of microbial metabolism. Nat Ecol Evol 5, 1424 (2021).10.1038/s41559-021-01535-8

9. V. Dubinkina, Y. Fridman, P. P. Pandey, S. Maslov, Multistability and regime shifts in microbial communities explained by competition for essential nutrients. Elife 8, e49720 (2019).10.7554/eLife.49720

10. C. Liao, T. Wang, S. Maslov, J. B. Xavier, Modeling microbial cross-feeding at intermediate scale portrays community dynamics and species coexistence. PLoS Comput Biol 16, e1008135 (2020).10.1371/journal.pcbi.1008135

11. P. Y. Ho, T. H. Nguyen, J. M. Sanchez, B. C. DeFelice, K. C. Huang, Resource competition predicts assembly of gut bacterial communities in vitro. Nat Microbiol 9, 1036 (2024).10.1038/s41564-024-01625-w

12. J. E. Goldford, N. Lu, D. Bajic, S. Estrela, M. Tikhonov, A. Sanchez-Gorostiaga, D. Segre, P. Mehta, A. Sanchez, Emergent simplicity in microbial community assembly. Science 361, 469 (2018).10.1126/science.aat1168

13. C. P. Mancuso, H. Lee, C. I. Abreu, J. Gore, A. S. Khalil, Environmental fluctuations reshape an unexpected diversity-disturbance relationship in a microbial community. Elife 10, (2021).10.7554/eLife.67175

14. D. Berry, S. Widder, Deciphering microbial interactions and detecting keystone species with co-occurrence networks. Front Microbiol 5, 219 (2014).10.3389/fmicb.2014.00219

15. D. Gonze, K. Z. Coyte, L. Lahti, K. Faust, Microbial communities as dynamical systems. Curr Opin Microbiol 44, 41 (2018).10.1016/j.mib.2018.07.004

16. X. Guo, J. Q. Boedicker, The Contribution of High-Order Metabolic Interactions to the Global Activity of a Four-Species Microbial Community. PLoS Comput Biol 12, e1005079 (2016).10.1371/journal.pcbi.1005079

17. M. S. Laska, J. T. Wootton, Theoretical concepts and empirical approaches to measuring interaction strength. Ecology 79, 461 (1998).DOI 10.1890/0012-9658(1998)079[0461:Tcaeat]2.0.Co;2

18. B. Momeni, L. Xie, W. Shou, Lotka-Volterra pairwise modeling fails to capture diverse pairwise microbial interactions. Elife 6, (2017).10.7554/eLife.25051

19. Y. Xiao, M. T. Angulo, J. Friedman, M. K. Waldor, S. T. Weiss, Y. Y. Liu, Mapping the ecological networks of microbial communities. Nat Commun 8, 2042 (2017).10.1038/s41467-017-02090-2

20. R. Marsland, 3rd, W. Cui, J. Goldford, A. Sanchez, K. Korolev, P. Mehta, Available energy fluxes drive a transition in the diversity, stability, and functional structure of microbial communities. PLoS Comput Biol 15, e1006793 (2019).10.1371/journal.pcbi.1006793

21. I. Guex, C. Mazza, M. Dubey, M. Batsch, R. Li, J. R. van der Meer, Regulated bacterial interaction networks: A mathematical framework to describe competitive growth under inclusion of metabolite cross-feeding. PLoS Comput Biol 19, e1011402 (2023).10.1371/journal.pcbi.1011402

22. O. Ponomarova, K. R. Patil, Metabolic interactions in microbial communities: untangling the Gordian knot. Curr Opin Microbiol 27, 37 (2015).10.1016/j.mib.2015.06.014

23. J. F. Yamagishi, N. Saito, K. Kaneko, Adaptation of metabolite leakiness leads to symbiotic chemical exchange and to a resilient microbial ecosystem. PLoS Comput Biol 17, e1009143 (2021).10.1371/journal.pcbi.1009143

24. N. Paczia, A. Nilgen, T. Lehmann, J. Gatgens, W. Wiechert, S. Noack, Extensive exometabolome analysis reveals extended overflow metabolism in various microorganisms. Microb Cell Fact 11, 122 (2012).10.1186/1475-2859-11-122

25. A. R. Pacheco, M. Moel, D. Segre, Costless metabolic secretions as drivers of interspecies interactions in microbial ecosystems. Nat Commun 10, 103 (2019).10.1038/s41467-018-07946-9

26. A. R. Pacheco, M. L. Osborne, D. Segre, Non-additive microbial community responses to environmental complexity. Nat Commun 12, 2365 (2021).10.1038/s41467-021-22426-3

27. T. Wang, A. Goyal, V. Dubinkina, S. Maslov, Evidence for a multi-level trophic organization of the human gut microbiome. PLoS Comput Biol 15, e1007524 (2019).10.1371/journal.pcbi.1007524

28. M. Schäfer, A. R. Pacheco, R. Kunzler, M. Bortfeld-Miller, C. M. Field, E. Vayena, V. Hatzimanikatis, J. A. Vorholt, Metabolic interaction models recapitulate leaf microbiota ecology. Science 381, eadf5121 (2023).10.1126/science.adf5121

29. L. Oña, S. Giri, N. Avermann, M. Kreienbaum, K. M. Thormann, C. Kost, Obligate cross-feeding expands the metabolic niche of bacteria. Nat Ecol Evol 5, 1224 (2021).10.1038/s41559-021-01505-0

30. D. W. Rivett, T. Bell, Abundance determines the functional role of bacterial phylotypes in complex communities. Nat Microbiol 3, 767 (2018).10.1038/s41564-018-0180-0

31. E. Bauer, J. Zimmermann, F. Baldini, I. Thiele, C. Kaleta, BacArena: Individual-based metabolic modeling of heterogeneous microbes in complex communities. PLoS Comput Biol 13, e1005544 (2017).10.1371/journal.pcbi.1005544

32. A. Dal Co, S. van Vliet, D. J. Kiviet, S. Schlegel, M. Ackermann, Short-range interactions govern the dynamics and functions of microbial communities. Nat Ecol Evol 4, 366 (2020).10.1038/s41559-019-1080-2

33. S. Čaušević, J. Tackmann, V. Sentchilo, L. Malfertheiner, C. von Mering, J. R. van der Meer, Habitat filtering more than microbiota origin controls microbiome transplant outcomes in soil. ISME J 19, (2025).10.1093/ismejo/wraf162

34. S. Čaušević, J. Tackmann, V. Sentchilo, C. von Mering, J. R. van der Meer, Reproducible propagation of species-rich soil bacterial communities suggests robust underlying deterministic principles of community formation. mSystems, e0016022 (2022).10.1128/msystems.00160-22

35. D. Hoces, G. Greter, M. Arnoldini, M. L. Stäubli, C. Moresi, A. Sintsova, S. Berent, I. Kolinko, F. Bansept, A. Woller, J. Hafliger, E. Martens, W. D. Hardt, S. Sunagawa, C. Loverdo, E. Slack, Fitness advantage of Bacteroides thetaiotaomicron capsular polysaccharide in the mouse gut depends on the resident microbiota. Elife 12, (2023).10.7554/eLife.81212

36. T. A. Joseph, P. Chlenski, A. Litman, T. Korem, I. Pe’er, Accurate and robust inference of microbial growth dynamics from metagenomic sequencing reveals personalized growth rates. Genome Res 32, 558 (2022).10.1101/gr.275533.121

37. Z. Pasternak, S. Pietrokovski, O. Rotem, U. Gophna, M. N. Lurie-Weinberger, E. Jurkevitch, By their genes ye shall know them: genomic signatures of predatory bacteria. ISME J 7, 756 (2013).10.1038/ismej.2012.149

38. H. Song, Y. Zhu, Z. Qu, M. Zhu, X. Li, L. Zhao, K. Wang, R. Zhang, L. Cui, Y. Li, Z. Bian, W. Zhang, Y. Chen, L. Du, J. L. Wang, X. Zhao, L. Deng, Y. Wang, Two-step localization driven by peptidoglycan hydrolase in interbacterial predation. ISME J 19, (2025).10.1093/ismejo/wraf208

39. J. D. Palmer, K. R. Foster, Bacterial species rarely work together. Science 376, 581 (2022).10.1126/science.abn5093

40. J. Kehe, A. Ortiz, A. Kulesa, J. Gore, P. C. Blainey, J. Friedman, Positive interactions are common among culturable bacteria. Sci Adv 7, eabi7159 (2021).10.1126/sciadv.abi7159

41. A. R. Pacheco, C. Pauvert, D. Kishore, D. Segre, Toward FAIR Representations of Microbial Interactions. mSystems 7, e0065922 (2022).10.1128/msystems.00659-22

42. A. Baichman-Kass, T. Song, J. Friedman, Competitive interactions between culturable bacteria are highly non-additive. Elife 12, (2023).10.7554/eLife.83398

43. L. S. Bittleston, M. Gralka, G. E. Leventhal, I. Mizrahi, O. X. Cordero, Context-dependent dynamics lead to the assembly of functionally distinct microbial communities. Nat Commun 11, 1440 (2020).10.1038/s41467-020-15169-0

44. M. Dubey, N. Hadadi, S. Pelet, N. Carraro, D. R. Johnson, J. R. van der Meer, Environmental connectivity controls diversity in soil microbial communities. Commun Biol 4, 492 (2021).10.1038/s42003-021-02023-2

45. T. N. Enke, M. S. Datta, J. Schwartzman, N. Cermak, D. Schmitz, J. Barrere, A. Pascual-Garcia, O. X. Cordero, Modular Assembly of Polysaccharide-Degrading Marine Microbial Communities. Curr Biol 29, 1528 (2019).10.1016/j.cub.2019.03.047

46. T. Miguel Trabajo, I. Guex, M. Dubey, E. Sarton-Loheac, H. Todorov, X. Richard, C. Mazza, J. R. van der Meer, Inferring bacterial interspecific interactions from microcolony growth expansion. Microlife 5, uqae020 (2024).10.1093/femsml/uqae020

47. V. Pandey, N. Hadadi, V. Hatzimanikatis, Enhanced flux prediction by integrating relative expression and relative metabolite abundance into thermodynamically consistent metabolic models. PLoS Comput Biol 15, e1007036 (2019).10.1371/journal.pcbi.1007036

48. O. Oftadeh, V. Hatzimanikatis, Genome-scale models of metabolism and expression predict the metabolic burden of recombinant protein expression. Metab Eng 84, 109 (2024).10.1016/j.ymben.2024.06.005

49. M. Vihola, Robust adaptive Metropolis algorithm with coerced acceptance rate. Stat Comput 22, 997 (2012).10.1007/s11222-011-9269-5

50. P. Servais, G. Billen, J. V. Rego, Rate of bacterial mortality in aquatic environments. Appl Environ Microbiol 49, 1448 (1985).10.1128/aem.49.6.1448-1454.1985

51. M. Loferer-Krossbacher, J. Klima, R. Psenner, Determination of bacterial cell dry mass by transmission electron microscopy and densitometric image analysis. Appl Environ Microbiol 64, 688 (1998)

52. D. J. Reasoner, E. E. Geldreich, A new medium for the enumeration and subculture of bacteria from potable water. Appl Environ Microbiol 49, 1 (1985)

53. S. Čaušević, M. Dubey, M. Morales, G. Salazar, V. Sentchilo, N. Carraro, H. J. Rüscheweyh, S. Sunagawa, J. R. van der Meer, Niche availability and competitive loss by facilitation control proliferation of bacterial strains intended for soil microbiome interventions. Nat Commun 15, 2557 (2024).10.1038/s41467-024-46933-1

54. S. Andrews, FastQC: A quality control tool for high throughput sequence data. (2015) https://www.bioinformatics.babraham.ac.uk/projects/fastqc/.

55. A. M. Bolger, M. Lohse, B. Usadel, Trimmomatic: a flexible trimmer for Illumina sequence data. Bioinformatics 30, 2114 (2014).10.1093/bioinformatics/btu170

56. T. Magoc, S. L. Salzberg, FLASH: fast length adjustment of short reads to improve genome assemblies. Bioinformatics 27, 2957 (2011).10.1093/bioinformatics/btr507

57. T. Lassmann, Kalign 3: multiple sequence alignment of large datasets. Bioinformatics 36, 1928 (2020). 10.1093/bioinformatics/btz795

58. B. Bushnell. (2014),

59. B. Langmead, S. L. Salzberg, Fast gapped-read alignment with Bowtie 2. Nat Methods 9, 357 (2012).10.1038/nmeth.1923

60. Y. Liao, G. K. Smyth, W. Shi, featureCounts: an efficient general purpose program for assigning sequence reads to genomic features. Bioinformatics 30, 923 (2014).10.1093/bioinformatics/btt656

61. H. Klingenberg, P. Meinicke, How to normalize metatranscriptomic count data for differential expression analysis. PeerJ 5, e3859 (2017).10.7717/peerj.3859

62. B. Muzellec, M. Telenczuk, V. Cabeli, M. Andreux, PyDESeq2: a python package for bulk RNA-seq differential expression analysis. Bioinformatics 39, (2023).10.1093/bioinformatics/btad547

63. C. P. Cantalapiedra, A. Hernandez-Plaza, I. Letunic, P. Bork, J. Huerta-Cepas, eggNOG-mapper v2: Functional Annotation, Orthology Assignments, and Domain Prediction at the Metagenomic Scale. Mol Biol Evol 38, 5825 (2021).10.1093/molbev/msab293

64. J. Zimmermann, C. Kaleta, S. Waschina, gapseq: informed prediction of bacterial metabolic pathways and reconstruction of accurate metabolic models. Genome Biol 22, 81 (2021).10.1186/s13059-021-02295-1

65. M. Kanehisa, Y. Sato, M. Kawashima, M. Furumichi, M. Tanabe, KEGG as a reference resource for gene and protein annotation. Nucleic Acids Res 44, D457 (2016).10.1093/nar/gkv1070

66. Y. Darzi, I. Letunic, P. Bork, T. Yamada, iPath3.0: interactive pathways explorer v3. Nucleic Acids Res 46, W510 (2018).10.1093/nar/gky299

67. M. Batsch, I. Guex, H. Todorov, C. M. Heiman, J. Vacheron, J. A. Vorholt, C. Keel, J. R. van der Meer, Fragmented micro-growth habitats present opportunities for alternative competitive outcomes. Nat Commun 15, 7591 (2024).10.1038/s41467-024-51944-z

