## Supplementary information for "Biphasic bacterial community assembly predicted from generalized first principles of monoculture growth and inferred species interactions"

table S1: Characteristics of the 21 soil isolates.

table S2: HILIC analysis of soil water extractable compounds.

Supplementary FIG 1. Simulation outcomes of soil-growth of the 20-member defined community with random SMINT-values.

Supplementary FIG 2. Community demography in the 21-member SynCom used for meta-transcriptomic analysis

Supplementary FIG 3. Prediction of population growth in the SynCom-21 based on CoPTR calculations.

Supplementary FIG 4. Classification of transporter and KEGG-orthology pathway expression types in the 21-membered SynCom used for metatranscriptomic analysis.

Supplementary FIG 5. Predicted substrate detail for any of the Type 1 (Peak at timepoint 2 or 3) or Type 2 (troughs) transporter expression profiles

Supplementary FIG 6. Principal component analysis comparison of ko-pathway annotated expression changes among the eight abundant species in the SynCom-21 metatranscriptomics experiment.

Supplementary FIG 7. Temporal changes in KEGG orthology metabolic pathways in the growing SynCom-21 metatranscriptomics experiment

Supplementary FIG 8. Predictions from the SynCom-21 SMINT/PARINT-model for growth of subcommunities with reduced composition.

Supplementary FIG 9. Predicted (A) and observed (B) community compositions of a weekly soil-diluted SynCom-21.

Supplementary FIG 10. Simulated effect of differences in inoculum composition on community development.

Supplementary FIG 11. Changing species interactions during SynCom-21 growth in liquid suspension.

Supplementary table 1. Characteristics of the 21 soil isolates.

| **Closest Species name** | **Strain number in collection** | **Abbre-viation** | **Estimated dry weight (fg C)** | **SynCom compositions** [a] | | | | | |
|---|---|---|---|---|---|---|---|---|---|
| | | | | **SynCom-20** | **SynCom-21** | **SynCom-21 MT** | **Sub-Com 1** | **Sub-Com 2** | **Sub-Com 3** |
| *Bradyrhizobium betae* | 6478 | *Bra* | *250* | green | green | green | salmon | salmon | salmon |
| *Burkholderia sp.* OLGA172 | 6699 | *Bur* | *290* | green | green | green | green | green | green |
| *Caulobacter sp.* Ji-3-8 | 6537 | *Cau* | *275* | green | green | green | green | green | green |
| *Cellulomonas xylanilytica* | 6483 | *Cel* | *250* | green | green | green | green | green | green |
| *Chitinophaga pinensis* | 6718 | *Chi* | *500* | green | green | green | green | green | green |
| *Cohnella sp.* MFER-1 | 6695 | *Coh* | *250* | green | green | green | salmon | salmon | green |
| *Curtobacterium pusillum* | 6155 | *Cur* | *250* | green | green | green | green | green | green |
| *Devosia riboflavina* | 6593 | *Dev* | *250* | green | green | green | green | green | green |
| *Flavobacterium sp.* KBS0721 | 6147 | *Fla* | *250* | green | green | green | green | green | green |
| *Luteibacter rhizovicinus* | 6500 | *Lut* | *250* | green | green | green | green | green | green |
| *Lysobacter gummosus* | 6525 | *Lys* | *275* | salmon | green | green | green | green | salmon |
| *Mesorhizobium loti* | 6519 | *Mes* | *250* | green | green | green | green | green | green |
| *Microbacterium sp.* PAMC 28756 | 6150 | *Mic* | *250* | green | green | green | green | green | green |
| *Mucilaginibacter ginsenosidivorax* | 6152 | *Muc* | *250* | green | green | green | salmon | salmon | salmon |
| *Phenylobacterium zucineum* | 6508 | *Phe* | *250* | green | green | green | salmon | salmon | salmon |
| *Pseudomonas moraviensis* | 6551 | *Pse-1* | *290* | green | green | green | green | salmon | green |
| *Pseudomonas sp.* CFSAN084952 | 6533 | *Pse-2* | *290* | green | green | green | green | green | green |
| *Rahnella sp.* CDC21234 | 6700 | *Rah* | *275* | green | green | green | salmon | salmon | salmon |
| *Rhodococcus fascians* | 6477 | *Rho* | *250* | green | green | green | green | green | green |
| *Tardiphaga robiniae* | 6697 | *Tar* | *250* | green | green | green | green | green | green |
| *Variovorax paradoxus* | 6146 | *Var* | *275* | green | green | green | green | green | green |

a) green = present, salmon = absent

Supplementary table 2: HILIC-MS/MS detected compounds in aqueous extract of soil microcosms inoculated with the 21-member community of soil isolates.

| Compounds | sterile control | SynCom Day1 | SynCom Day3 | SynCom Day10 |
|---|---|---|---|---|
| 1-METHYLADENOSINE | 2.74E+05[a] | 4.72E+03 | 9.67E-04 | 1.04E-03 |
| 2-HYDROXYBUTYRATE | 8.38E+03 | 5.02E+03 | 3.56E+02 | 4.43E+02 |
| 2-HYDROXYGLUTARIC ACID | 7.69E+03 | 2.44E+03 | 1.09E-03 | 1.08E+02 |
| 3-(2-HYDROXYPHENYL)PROPANOATE | 1.22E+05 | 3.07E+04 | 6.07E+03 | 4.07E+03 |
| 3-HYDROXYBUTANOATE | 1.34E+04 | 1.62E+04 | 7.50E+02 | 5.54E+02 |
| 3-HYDROXYBUTYRATE | 4.47E+04 | 1.98E+02 | 9.90E-04 | 9.98E-04 |
| 3-METHOXY-4-HYDROXYMANDELATE | 8.98E+03 | 7.92E+03 | 8.55E-04 | 1.26E-03 |
| 4-ACETAMIDOBUTANOATE | 4.46E+03 | 4.40E+03 | 8.68E+03 | 5.27E+03 |
| 4-GUANIDINOBUTANOATE | 4.24E+05 | 6.91E+04 | 7.21E+04 | 8.59E+04 |
| 4-HYDROXYBENZOATE | 6.35E+04 | 1.27E+03 | 1.13E+03 | 1.37E+03 |
| 4-IMIDAZOLEACETATE | 6.13E+04 | 5.96E+04 | 5.89E+04 | 5.01E+04 |
| 4-PYRIDOXATE | 4.34E+04 | 4.72E+04 | 1.26E+05 | 1.36E+05 |
| 5'-DEOXYADENOSINE | 4.26E+04 | 9.27E+03 | 7.52E+03 | 6.35E+03 |
| 5-HYDROXYINDOLEACETATE | 3.73E+03 | 3.43E+03 | 3.71E+03 | 3.06E+03 |
| 5-HYDROXYTRYPTOPHAN | 4.97E+03 | 7.79E+03 | 5.72E+03 | 4.87E+03 |
| 5-METHYLCYTOSINE | 1.51E+03 | 1.89E+03 | 1.43E+03 | 7.55E+02 |
| 6-HYDROXYMELATONIN | 1.41E+03 | 1.37E+03 | 1.38E+03 | 1.20E+03 |
| ACETYLCARNITINE | 9.25E+04 | 7.04E+04 | 4.77E+03 | 4.22E+03 |
| ADENOSINE | 2.34E+07 | 5.99E+04 | 4.41E+04 | 3.89E+04 |
| ALANINE | 1.27E+05 | 1.05E-03 | 1.03E-03 | 1.03E-03 |
| ALLANTOIN | 3.22E+03 | 1.65E+02 | 5.37E+01 | 1.03E-03 |
| AMP | 5.40E+03 | 2.51E+03 | 2.52E+02 | 1.21E-03 |
| ARGININE | 2.25E+04 | 7.07E+03 | 1.12E+04 | 9.95E+03 |
| ASPARAGINE | 4.88E+03 | 5.94E+02 | 9.60E-04 | 9.99E-04 |
| ASPARTATE | 2.89E+04 | 4.02E+03 | 6.00E+02 | 3.18E+02 |
| BETA-ALANINE/SARCOSINE | 2.05E+04 | 5.61E+01 | 9.37E+01 | 9.98E-04 |
| BETAINE | 1.53E+07 | 1.02E-03 | 1.04E-03 | 9.73E-04 |
| BUTYRYLCARNITINE | 6.42E+04 | 4.87E+04 | 2.83E+04 | 2.04E+04 |
| CARNITINE | 3.26E+05 | 5.77E+03 | 5.43E+03 | 6.15E+03 |
| CHOLINE | 6.92E+06 | 5.27E+04 | 4.07E+04 | 9.41E+04 |
| CIS-ACONITATE | 9.04E+04 | 4.91E+03 | 4.11E+03 | 4.55E+03 |
| CITRATE_1 | 2.96E+05 | 2.48E+05 | 2.27E+05 | 2.71E+05 |
| CITRULLINE | 2.91E+04 | 7.64E+03 | 6.03E+03 | 4.43E+03 |
| CREATINE | 1.52E+04 | 1.22E+04 | 1.30E+04 | 1.21E+04 |
| CREATININE | 1.37E+05 | 1.22E+05 | 9.99E+04 | 8.54E+04 |
| CYCLIC GMP | 6.20E+02 | 8.24E+01 | 2.67E+02 | 1.01E+02 |
| CYTIDINE | 2.00E+05 | 1.51E+03 | 4.37E+02 | 6.35E+02 |

| | | | | |
|---|---|---|---|---|
| DECANOYLCARNITINE | 5.12E+03 | 7.91E+03 | 4.81E+03 | 6.99E+03 |
| DEOXYADENOSINE | 4.40E+04 | 1.01E+04 | 1.02E+04 | 5.99E+03 |
| DEOXYCARNITINE | 9.69E+05 | 1.45E+05 | 1.44E+05 | 1.85E+05 |
| DEOXYCYTIDINE | 2.23E+02 | 1.02E-03 | 2.25E+02 | 3.82E+02 |
| DEOXYURIDINE | 6.88E+03 | 1.03E-03 | 3.25E+02 | 1.24E+02 |
| DIETHANOLAMINE | 2.65E+04 | 2.67E+04 | 3.65E+04 | 4.78E+04 |
| DIHYDROURACIL | 9.54E+02 | 1.90E+03 | 2.26E+03 | 1.75E+03 |
| DIMETHYLGLYCINE | 2.78E+04 | 9.80E-04 | 9.70E-04 | 1.01E-03 |
| ETHANOLAMINE | 9.52E+04 | 2.45E+03 | 2.38E+03 | 2.35E+03 |
| GLUCONATE | 1.28E+05 | 8.39E+04 | 1.97E+03 | 1.54E+03 |
| GLUTAMATE | 9.20E+04 | 1.67E+03 | 1.14E+03 | 1.09E+03 |
| GLUTAMINE | 9.84E+03 | 1.43E+03 | 1.00E+03 | 5.51E+02 |
| GLYCERALDEHYDE 3-PHOSPHATE | 2.82E+03 | 2.36E+03 | 3.03E+03 | 2.62E+03 |
| GLYCERATE | 6.92E+04 | 1.62E+04 | 4.21E+02 | 5.87E+02 |
| GLYCEROL | 9.59E+04 | 1.14E+05 | 1.40E+05 | 1.42E+05 |
| GLYCEROL 3-PHOSPHATE | 2.08E+04 | 1.57E+03 | 1.20E+03 | 8.06E+02 |
| GUAIACOL | 1.32E+03 | 1.02E+03 | 1.79E+03 | 1.62E+03 |
| GUANINE | 1.99E+05 | 2.29E+03 | 1.74E+03 | 1.45E+03 |
| GUANOSINE | 1.01E+05 | 7.89E+02 | 3.34E+02 | 2.04E+02 |
| GUANOSINE MONOPHOSPHATE | 4.83E+03 | 2.38E+03 | 2.80E+02 | 5.63E+01 |
| GULONOLACTONE | 2.78E+03 | 3.61E+03 | 4.25E+03 | 4.04E+03 |
| HEXANOYLCARNITINE | 1.95E+03 | 1.79E+03 | 1.61E+03 | 1.64E+03 |
| HISTAMINE | 7.66E+02 | 8.44E+02 | 1.01E+03 | 1.51E+03 |
| HISTIDINE | 3.16E+04 | 2.30E+04 | 2.76E+04 | 2.25E+04 |
| HOMOGENTISATE | 2.26E+04 | 2.64E+04 | 1.40E+02 | 5.39E+01 |
| HYDROXYPHENYLLACTATE | 5.88E+03 | 4.27E+03 | 8.67E-04 | 1.23E-03 |
| HYPOXANTHINE | 1.88E+03 | 7.50E+01 | 8.67E+01 | 6.87E+01 |
| INDOLE-3-ACETAMIDE | 3.25E+02 | 3.47E+02 | 4.03E+02 | 6.73E+02 |
| INOSINE | 1.42E+04 | 9.97E-04 | 1.23E+02 | 1.06E-03 |
| ISOCITRATE | 4.75E+03 | 2.14E+03 | 2.30E+03 | 2.84E+03 |
| ISOLEUCINE | 2.24E+05 | 4.32E+03 | 1.00E-03 | 1.02E-03 |
| ISOVALERYLCARNITINE | 2.86E+04 | 2.35E+04 | 2.03E+04 | 1.62E+04 |
| KYNURENATE | 3.64E+04 | 3.60E+04 | 6.20E+03 | 7.01E+03 |
| LACTATE | 6.60E+03 | 2.32E+03 | 1.36E+03 | 1.07E+03 |
| LEUCINE | 5.54E+05 | 2.01E+05 | 2.17E+05 | 2.36E+05 |
| MALATE | 7.25E+05 | 2.68E+04 | 5.92E+03 | 5.65E+03 |
| MELATONIN | 4.45E+04 | 4.65E+04 | 4.08E+04 | 4.10E+04 |
| METANEPHRINE | 3.11E+03 | 4.10E+03 | 5.19E+03 | 4.93E+03 |
| METHYLGUANIDINE | 3.90E+05 | 3.67E+05 | 4.31E+05 | 4.36E+05 |
| METHYLTHIOADENOSINE | 2.25E+04 | 4.25E+02 | 5.80E+02 | 1.02E+03 |
| MEVALONATE | 6.38E+04 | 4.85E+04 | 9.46E-04 | 1.08E-03 |
| N,N,N-TRIMETHYLLYSINE | 3.24E+03 | 6.35E+03 | 2.88E+04 | 2.29E+04 |

| | | | | |
|---|---|---|---|---|
| N-ACETYLALANINE | 1.48E+04 | 5.27E+03 | 8.18E+01 | 2.94E+02 |
| N-ACETYLGLUTAMATE | 1.24E+04 | 2.75E+03 | 9.69E+02 | 9.01E+02 |
| N-ACETYLLEUCINE | 1.49E+04 | 3.08E+04 | 4.50E+03 | 4.06E+03 |
| N-ACETYLPHENYLALANINE | 2.01E+03 | 1.13E+04 | 8.61E+02 | 8.07E+02 |
| N-ACETYLSERINE | 6.73E+03 | 7.38E+03 | 8.09E+03 | 9.89E+03 |
| N-ACETYLTRYPTOPHAN | 6.25E+03 | 7.85E+03 | 6.57E+03 | 5.65E+03 |
| N-ALPHA-ACETYLLYSINE | 3.50E+04 | 5.05E+04 | 1.86E+04 | 7.18E+03 |
| N-METHYLASPARTATE | 1.57E+04 | 8.23E+03 | 1.08E-03 | 9.60E-04 |
| NAD | 8.74E+00 | 9.36E+02 | 6.46E+01 | 2.74E+02 |
| NICOTINAMIDE | 1.81E+05 | 9.06E+03 | 1.41E+04 | 1.98E+04 |
| NICOTINATE | 9.87E+03 | 1.41E+04 | 2.40E+03 | 2.24E+03 |
| OROTATE | 1.38E+04 | 1.23E+03 | 1.65E+03 | 1.76E+03 |
| OXOPROLINE | 7.23E+05 | 5.00E+04 | 2.93E+04 | 2.49E+04 |
| PANTOTHENATE | 1.02E+04 | 2.22E+04 | 2.90E+03 | 2.81E+03 |
| PARAXANTHINE | 1.39E+03 | 1.29E+03 | 1.40E+03 | 1.06E+03 |
| PHENYLACETALDEHYDE | 4.72E+03 | 3.97E+03 | 6.93E+02 | 6.73E+02 |
| PHENYLALANINE | 5.76E+04 | 4.78E+03 | 4.30E+03 | 3.42E+03 |
| PIPECOLATE | 1.42E+04 | 4.38E+03 | 2.45E+03 | 2.67E+03 |
| PROLINE | 5.79E+05 | 1.29E+04 | 1.12E+04 | 1.03E+04 |
| PROPIOYLCARNITINE | 1.58E+04 | 1.22E+04 | 3.72E+03 | 1.57E+03 |
| PYRIDOXAMINE | 3.27E+04 | 3.24E+04 | 3.83E+04 | 3.73E+04 |
| PYRIDOXINE | 2.35E+02 | 3.56E+02 | 7.41E+02 | 6.04E+02 |
| PYRIMIDINE | 1.19E+05 | 9.74E+04 | 8.49E+04 | 6.39E+04 |
| PYROCATECHOL | 1.50E+03 | 1.32E+03 | 1.34E+03 | 1.14E+03 |
| PYRUVATE | 3.92E+04 | 3.78E+03 | 1.27E+03 | 1.56E+03 |
| QUINATE | 4.38E+05 | 3.34E+03 | 2.15E+03 | 2.01E+03 |
| QUINOLINATE | 7.25E+03 | 7.99E+03 | 8.80E+03 | 8.25E+03 |
| SALICYLATE | 1.01E+05 | 1.04E+05 | 9.73E+03 | 1.12E+04 |
| SERINE | 2.12E+04 | 2.50E+03 | 2.85E+03 | 1.92E+03 |
| SORBITOL/MANNITOL | 8.73E+05 | 1.31E+04 | 9.86E-04 | 2.43E+04 |
| THYMIDINE | 6.25E+03 | 1.08E-03 | 3.71E+02 | 1.00E-03 |
| TRIGONELLINE | 9.71E+05 | 5.86E+05 | 4.29E+05 | 5.69E+05 |
| TRYPTAMINE | 1.56E+02 | 1.32E+02 | 1.72E+02 | 1.62E+02 |
| TYRAMINE | 9.38E+03 | 1.68E+04 | 5.33E+04 | 2.60E+04 |
| TYROSINE | 4.13E+03 | 2.43E+02 | 2.30E+02 | 1.01E-03 |
| URACIL | 3.36E+04 | 6.52E+02 | 1.80E+03 | 2.06E+03 |
| URATE | 3.93E+03 | 1.53E+03 | 1.23E+04 | 1.49E+04 |
| URIDINE | 1.91E+05 | 1.77E+03 | 1.27E+03 | 1.08E+03 |
| URIDINE MONOPHOSPHATE | 5.99E+03 | 1.75E+03 | 2.64E+02 | 1.10E-03 |
| VALINE | 4.67E+04 | 9.76E-04 | 5.82E+02 | 9.92E-04 |
| XANTHINE | 5.96E+03 | 1.09E+03 | 8.45E+02 | 1.74E+03 |
| XANTHOSINE | 4.45E+03 | 1.71E+03 | 9.44E-04 | 1.13E-03 |

a) mean peak area -ion counts; of three biological replicates.

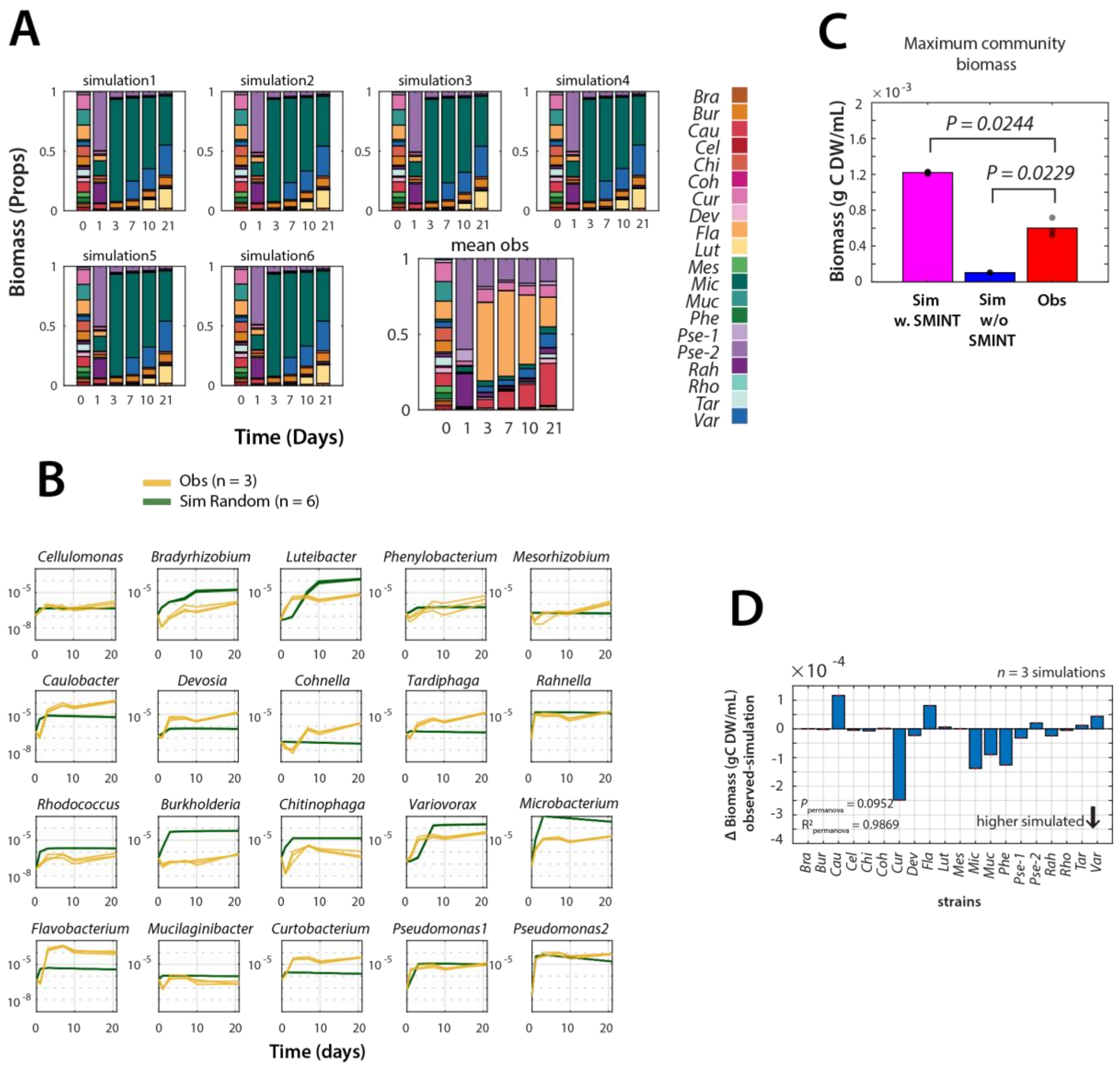


**Supplementary FIG 1. Simulation outcomes of soil-growth of the 20-member defined community with random SMINT-values. A)** Relative abundances of SynCom-20 strains over time in replicate simulations (*n* = 6) with SMINT-parameter values randomly sampled from the species' log-normal growth rate distributions multiplied by 0.8, in comparison to experimentally observed values, and **B)** their individual population growth profiles. **C)** Maximum observed (Obs) and simulated summed community biomass for the random starting conditions without (w/o) and with SMINT-optimized parameters. *P* values from a permutation test on the Cliff's delta index with n = 100,000 permutations. **D)** Net per species biomass difference at day 21 between simulated and observed conditions. *P* values derived from PERMANOVA using Bray–Curtis distances. $R^2$ represents the coefficient of determination, as explained in Fig. 3.

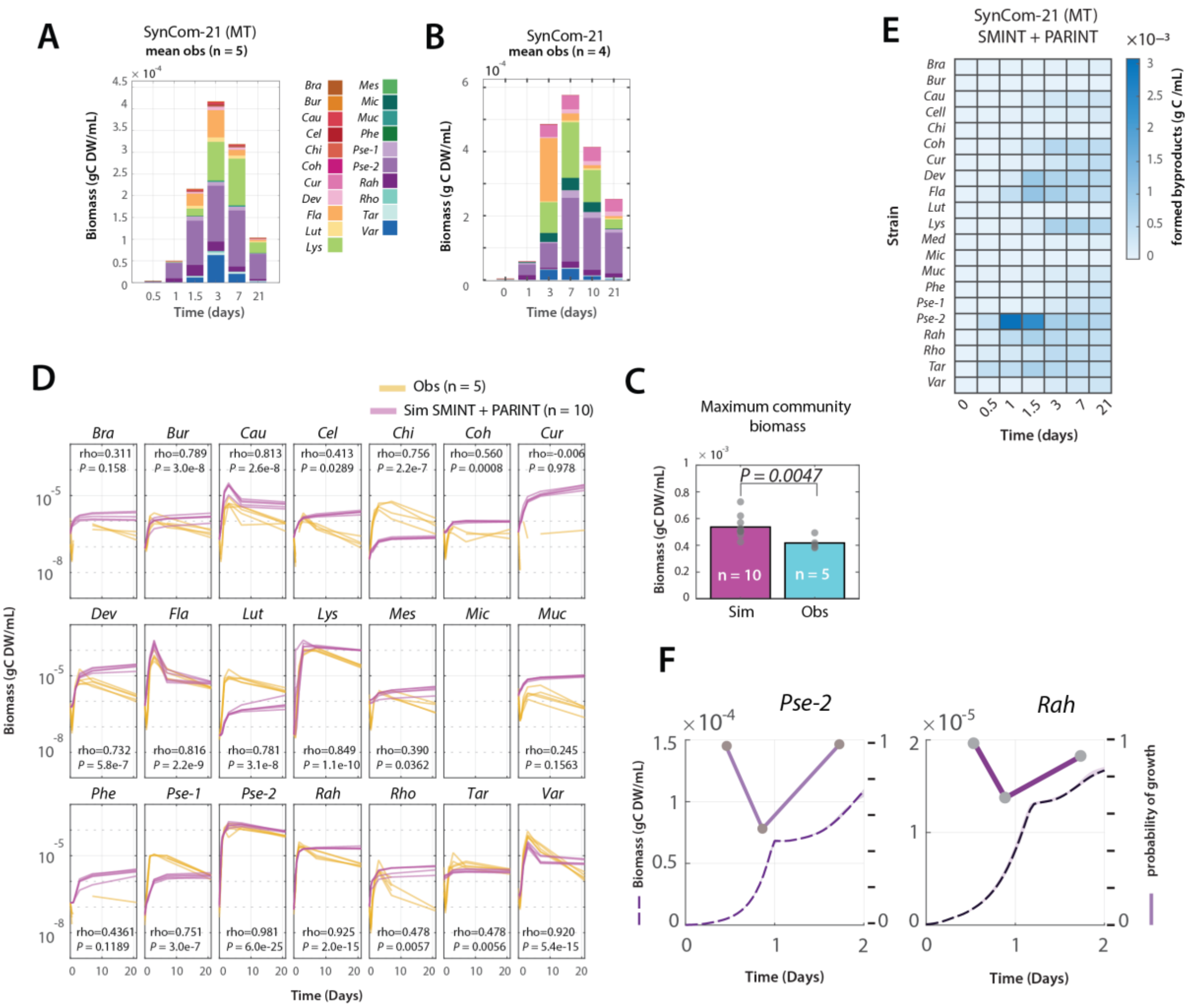


**Supplementary FIG 2. Community demography in the 21-member SynCom used for meta-transcriptomic analysis. A)** Observed absolute species abundances over time in five experimental soil microcosm replicates inoculated with the 21 soil isolates. MT, meta-transcriptomics. **B)** Reproduced main Fig. 4A of the SynCom-21 growth in the experiment used for optimization of the PARINT-parameters, for comparison to (A). **C)** Predicted (Sim, both SMINT- and PARINT-parameters) and observed maximum total biomass in the SynCom-21 MT experiment. *P* values from a permutation test on the Cliff's delta index with n = 100,000 permutations. **D)** Individual predicted and observed population growth of the 21 SynCom members, each in five replicates. rho and *P* correspond to the Pearson correlations between observed and predicted population trajectories (all replicates included individually). **E)** Predicted formed byproducts by each individual SynCom-21 member over time. **F)** Higher time resolution detail of the predicted biphasic growth of *Pseudomonas* sp. strain 2 and *Rahnella* in the SynCom-21 MT (dashed lines, both SMINT- and PARINT-parameters) and the corresponding measured probability of growth (solid lines) from meta-transcriptomic analysis (reproduced from main Fig. 6A).

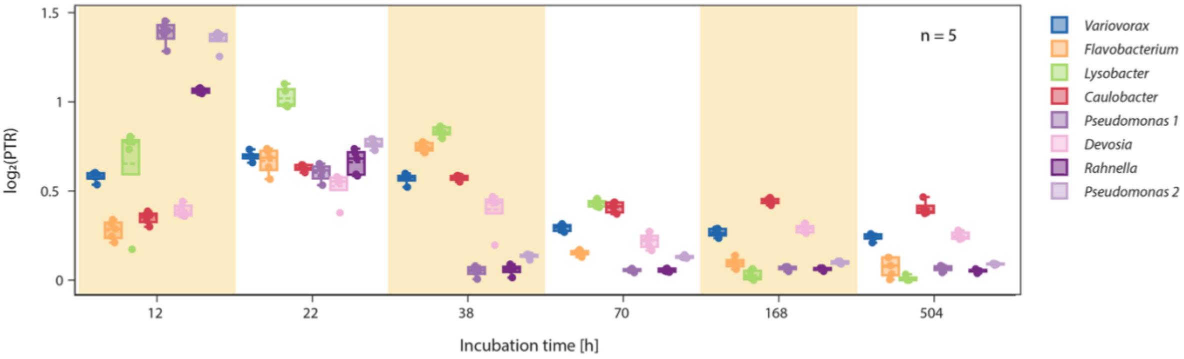


**Supplementary FIG 3. Prediction of population growth in the SynCom-21 based on CoPTR calculations.** Panels show the $\log_2$ 'peak-to-trough' ratio of the *origin* and terminus of chromosome replication regions quantified from metagenomic DNA isolated from the SynCom-21 recovered cell fractions for each of the listed SynCom-members shown on the right. Data are presented as box plots with overlaid individual data points from each of the replicate microcosms (n = 5).

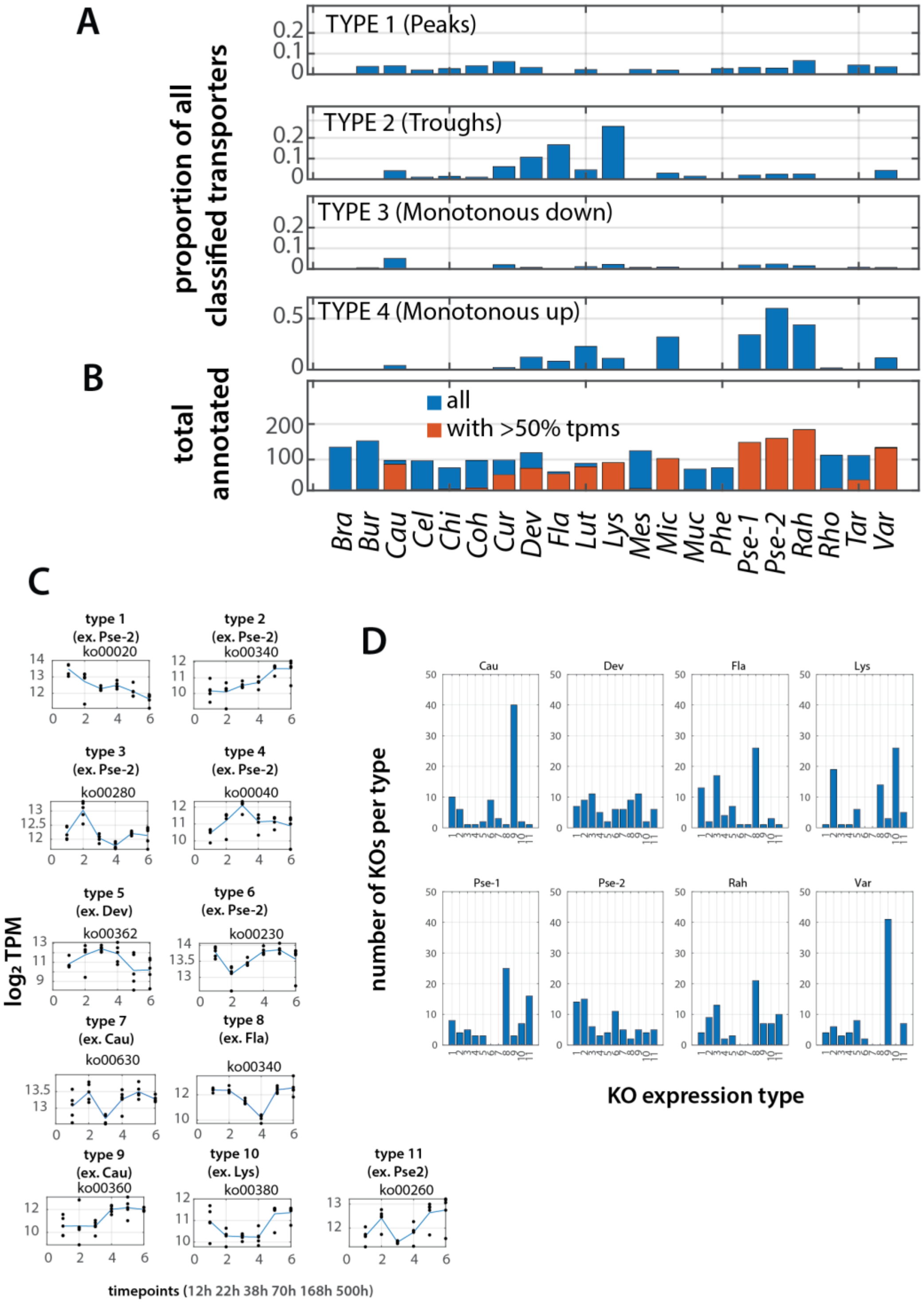


**Supplementary FIG 4. Classification of transporter and KEGG-orthology pathway expression types in the 21-membered SynCom used for metatranscriptomic analysis. A)** Proportion of classified transporters per genome with observed expression profiles over time showing a peak at timepoint 2 (22 h) or 3 (38 h; mean $log_2$ expression higher than any of the other timepoints + 2), a trough/valley at timepoint 2 or 3 (mean $log_2$ expression + 2 is lower than any of the other timepoints), monotonous increase (rho from Pearson correlation over all timepoints is > 0.5 and p-value < 0.05) or decrease (Pearson rho < -0.5) over time. Example expression profiles shown in main Fig. 6E. **B)** Total number of annotated transporters overlaid in red with the number of annotated transporters for those genomes with at least half of sample replicates and time points having detectable expression levels (transcripts per million, TPM). Only those being reported in main Fig. 6. **C)** Examples of categorized ko-pathway time expression profiles and **D)** the number of observed ko-pathway time expression profiles per genome and per category. Categorization based on maximum Pearson correlation to any of the example categories in (C).

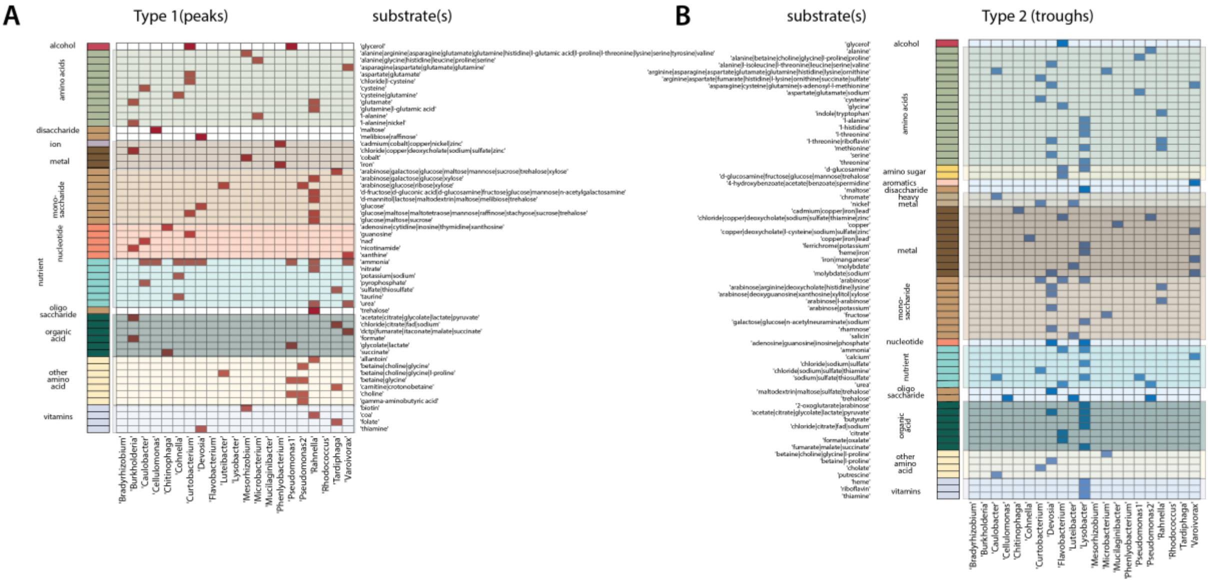


**Supplementary FIG 5. Predicted substrate detail for any of the Type 1 (Peak at timepoint 2 or 3) or Type 2 (troughs) transporter expression profiles. A)** Presence (red) or absence of classified transporter per genome and substrate group (on the left), and with predicted substrate(s) on the right. Detail for main Fig. 6G. **B)** as (A) but for transporter expression profiles of the Type 2 group.

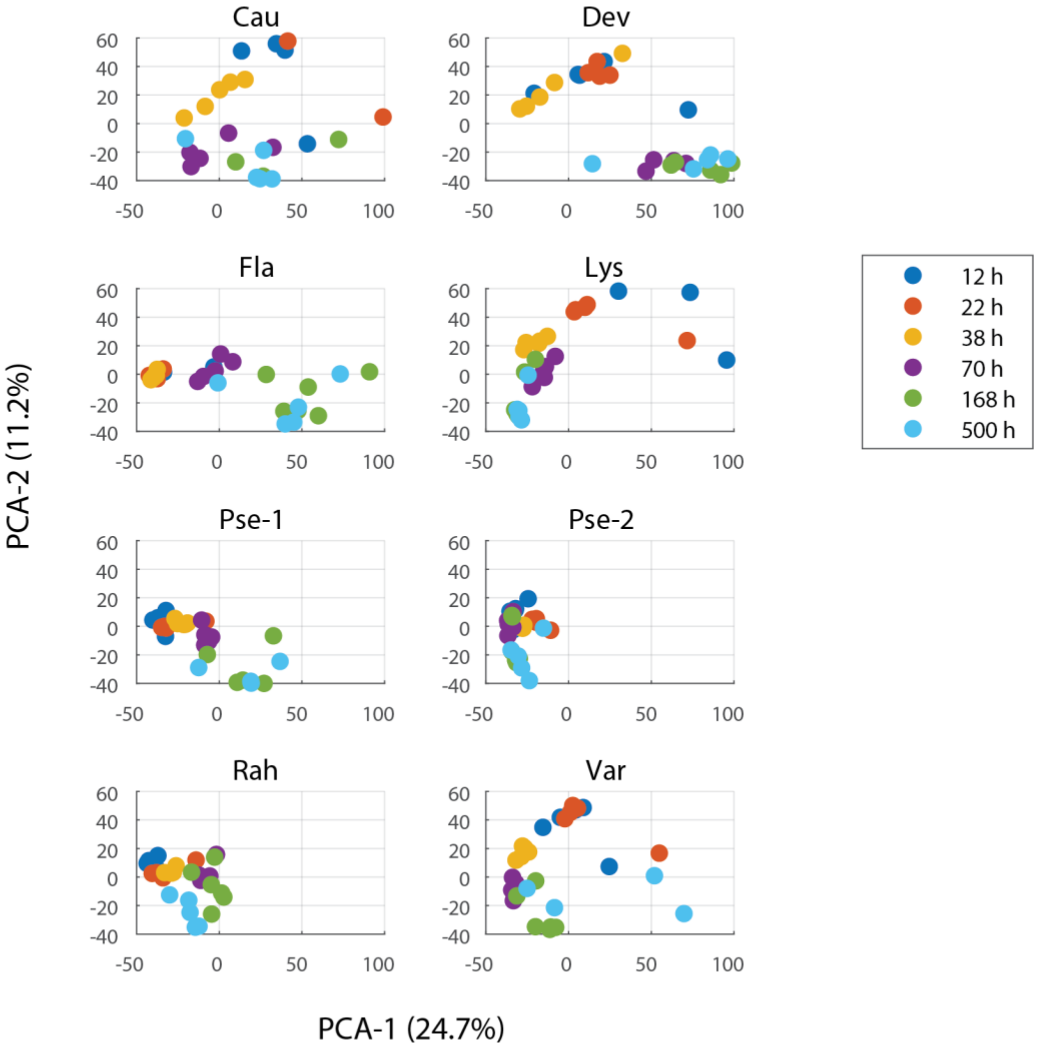


**Supplementary FIG 6. Principal component analysis comparison of ko-pathway annotated expression changes among the eight abundant species in the SynCom-21 metatranscriptomics experiment.** Panels show the two first principal component (PCA) coordinates in a combined PCA but separated by species for clarity. PCA based on expression values of genes annotated to common ko-pathway terms in order to compare across species. Circles show individual replicates colored by sampling time point in the developing SynCom-21. Percentages indicate the variation explained by the component. Species abbreviations: Cau, *Caulobacter*; Dev, *Devosia*; Fla, *Flavobacterium*; Lys, *Lysobacter*; Pse-1, *Pseudomonas sp.* strain 1; Pse-2, *Pseudomonas sp.* strain 2; Rah, *Rahnella*; Var, *Variovorax*.

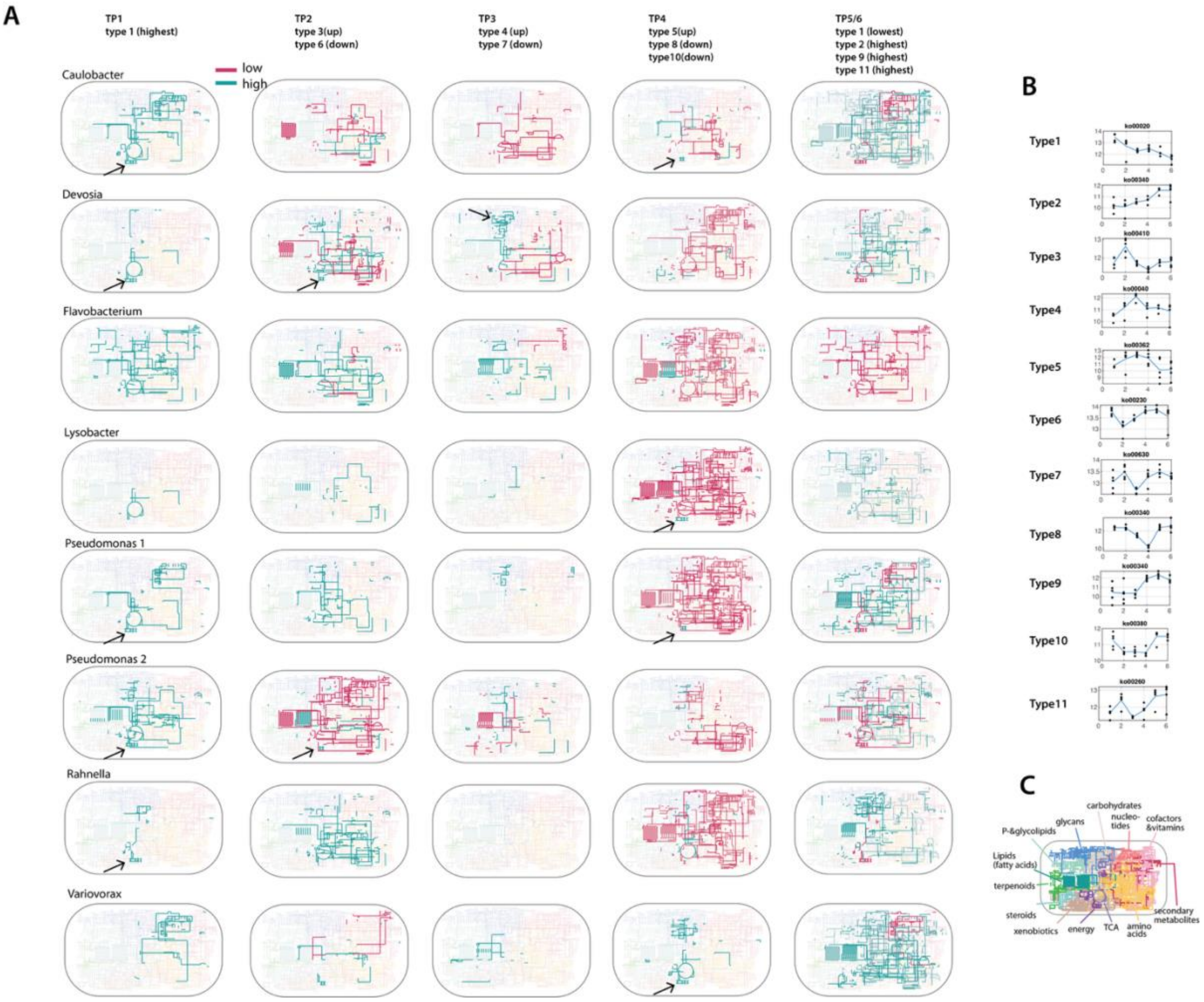


**Supplementary FIG 7. Temporal changes in KEGG orthology metabolic pathways in the growing SynCom-21 metatranscriptomics experiment. A)** Expression increase (blue) or decrease (red) in metabolic pathways grouped by expression types (in **B**) mapped on the global KO metabolic pathway overview of iPath3 (*1*) for eight of the most abundant SynCom-21 species with sufficient expression data available. Arrows point to examples of oxidative phosphorylation (*energy* metabolism) and *sugar* metabolism (*Devosia*). **B)** Repetition of the used KO-expression types that are summed for the timepoints (TP) in panel (A). **C)** Map location and used color (background in panel A maps) of the main KO pathways (reproduced from main Fig. 7B for clarity). TP, timepoints; TP1, 12h; TP2, 22h; TP3, 38h; TP4, 60h; TP5, 160 h; TP6, 500 h.

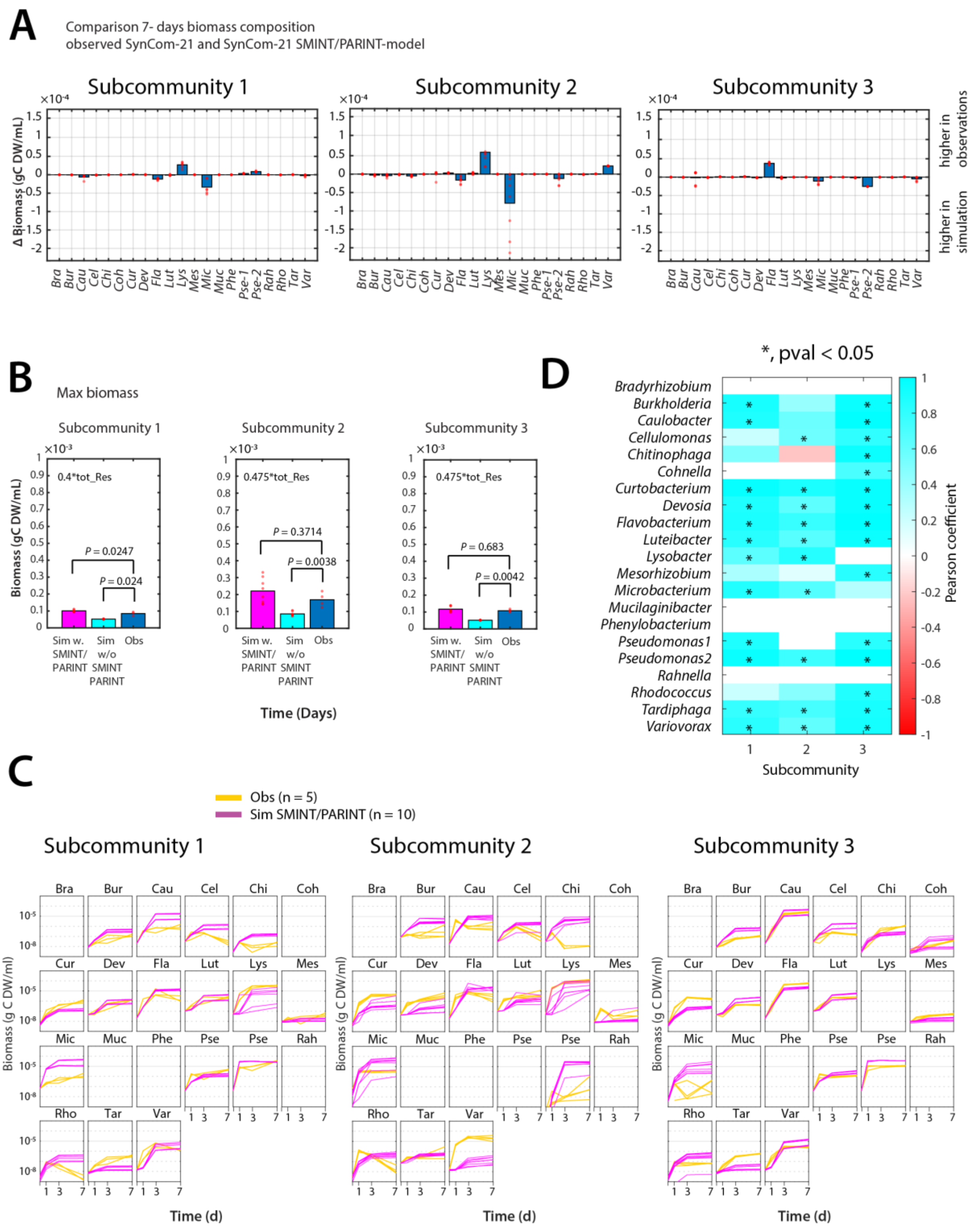


**Supplementary FIG 8. Predictions from the SynCom-21 SMINT/PARINT-model for growth of subcommunities with reduced composition. A)** Comparison of individual population biomass attained after day 7 in the experimental datasets and the simulations. **B)** Maximum attained biomass in the three subcommunities for the experimental datasets (obs), simulations with SMINT- and PARINT-parameters, or without (n = 10 simulations and 5 biological replicates). Resource concentration (tot_Res) in these experiments was less than in other soil microcosms (newly prepared soil extract with lower carbon mass). *P* values corresponds to permutation test on the Cliff's delta index with n = 100,000 permutations. **C)** Individual observed (yellow) and simulated (magenta) population growth, and their corresponding Pearson correlation coefficients (in **D**). Pearson correlation coefficients with *P* value < 0.05 are indicated with an asterisk.

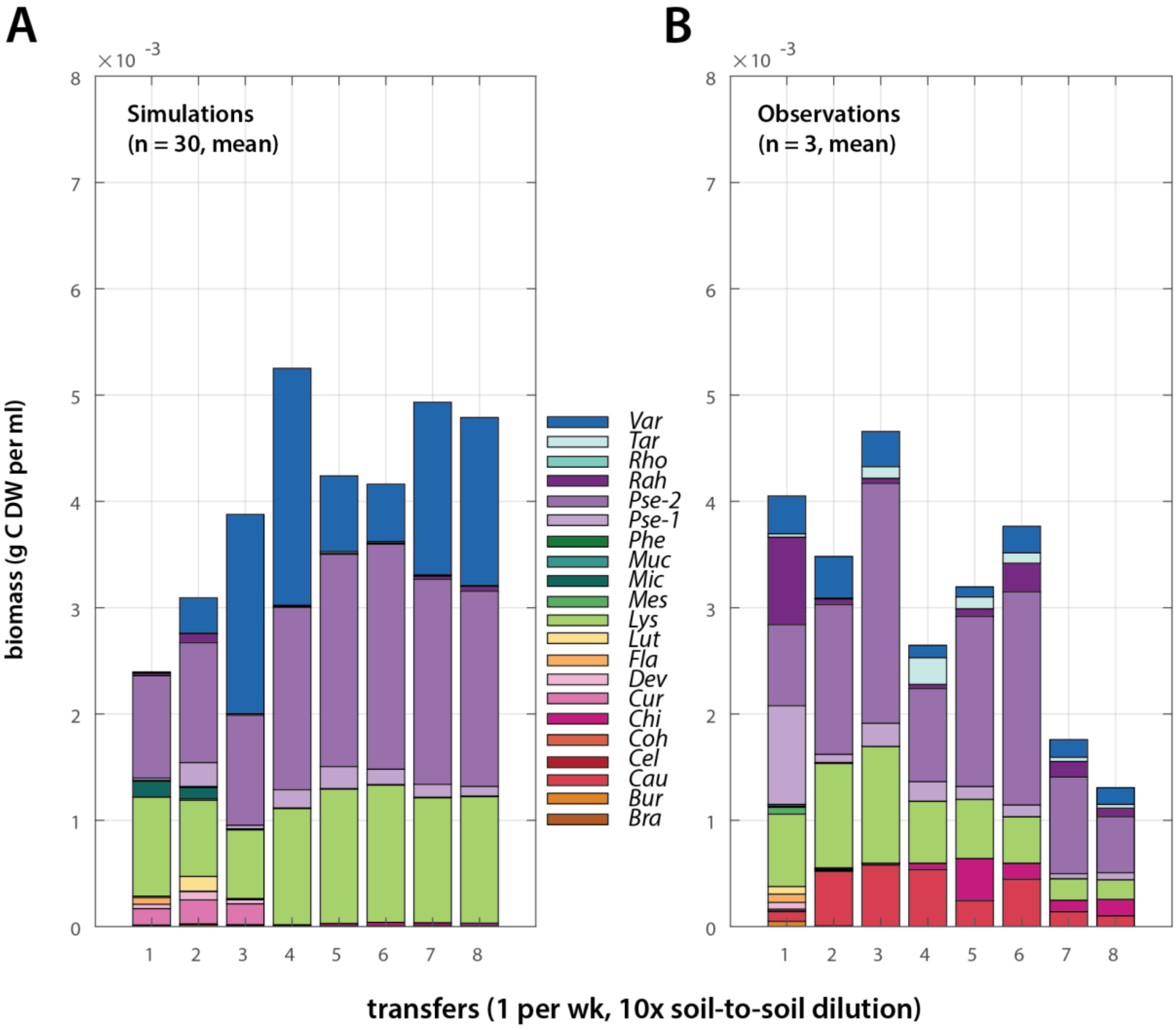


**Supplementary FIG 9. Predicted (A) and observed (B) community compositions of a weekly soil-diluted SynCom-21.** Stackplots represent absolute species abundances after 1 week incubation; color as per the legend. Soil-grown communities were diluted 1:10 in fresh sterile soil microcosms and again grown for 1 week. Eight transfers in total. Simulated community stackplots are the mean of 30 simulations that start with the experimentally observed composition after each transfer, but maintain the same sampled coefficient values from the SMINT/PARINT optimized parameters throughout. Experimental data taken from Ref. (*2*).

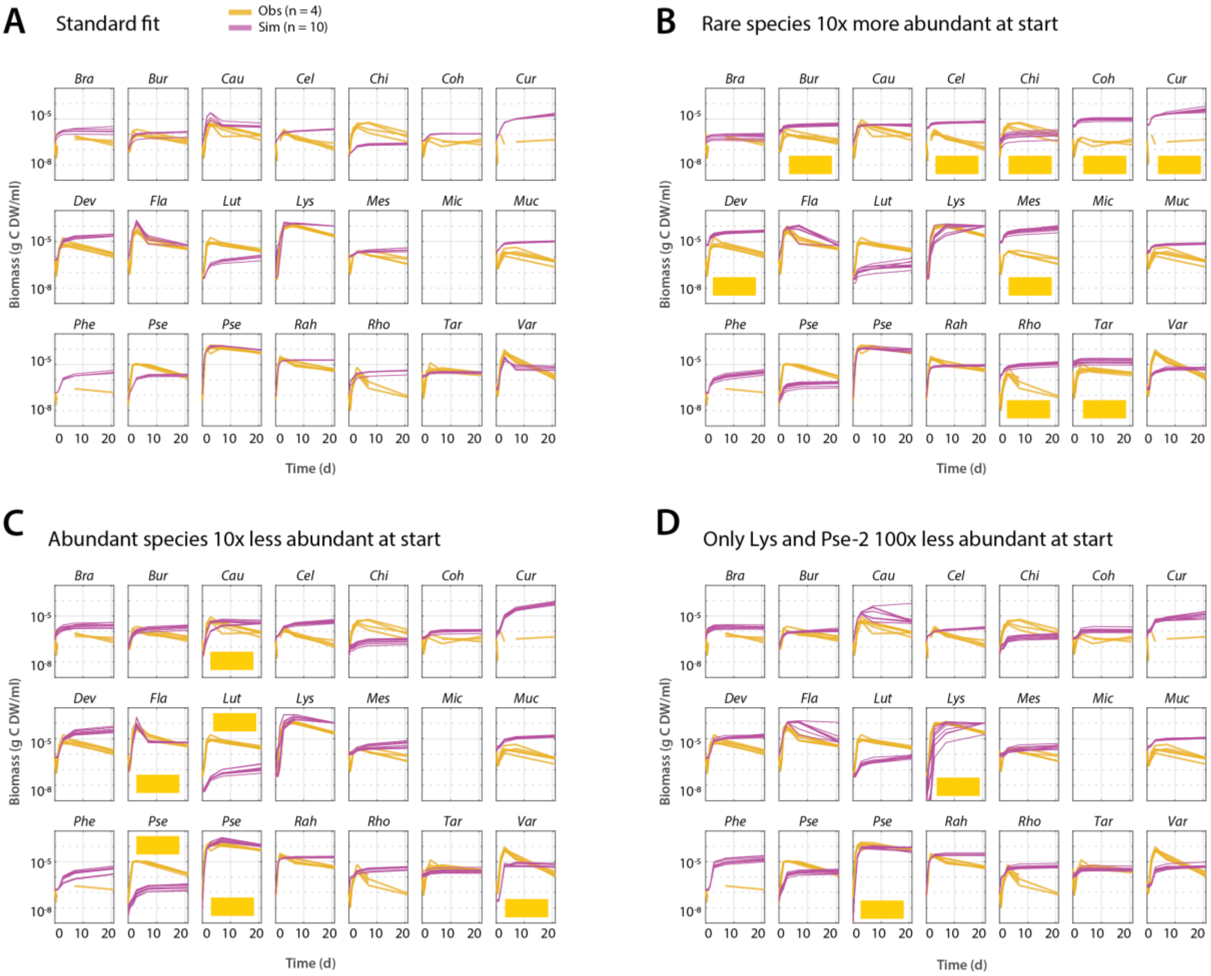


**Supplementary FIG 10. Simulated effect of differences in inoculum composition on community development.** Panels show individual simulated population growth (magenta, 10 replicates) in comparison to data from the SynCom-21 meta-transcriptomics experiment (blue, 4 replicates). **A)** Starting situation (reproduced from fig. S2C for clarity). **B)** Rare species abundances (marked in yellow) 10x higher at start. **C)** Abundant species (marked in yellow) 10x less abundant at start. **D)** Only *Lysobacter* and *Pseudomonas* sp. strain 2 100x less abundant at start.

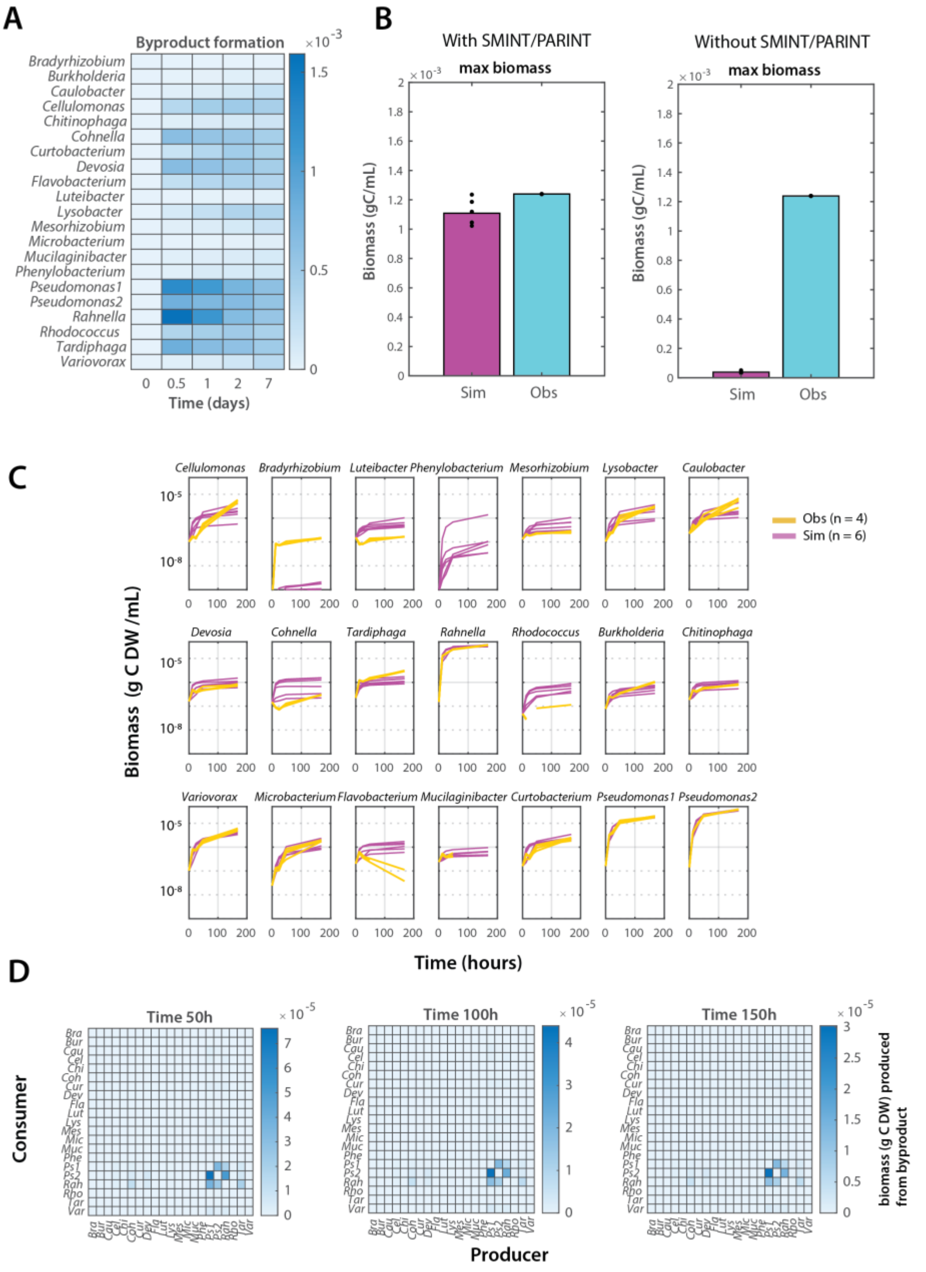


**Supplementary FIG 11. Changing species interactions during SynCom-21 growth in liquid suspension. A)** Byproduct formation calculated from optimized SMINT/PARINT parameters searched by SAO on data from a mixed liquid-grown SynCom-21 on soil extract. **B)** Observed and simulated biomass for the mixed liquid-grown SynCom-21 without and with its optimized SMINT-parameters. **C)** Individual observed population growth (yellow, *n* = 4 replicates) and trained simulated growth (magenta, *n* = 6 replicates). **D)** Predicted biomass formation from released byproducts (producer) and their consumption (consumer) at three time points using the the liquid-optimized SMINT- and PARINT-parameter datasets. Since this is a different growth habitat we did not use the soil-optimized SynCom-21 parameters, but specifically applied the SAO to the liquid experimental data.

**Supplementary references**


1. Y. Darzi, I. Letunic, P. Bork, T. Yamada, iPath3.0: interactive pathways explorer v3. *Nucleic Acids Res* **46**, W510 (2018).10.1093/nar/gky299
2. S. Čaušević, J. Tackmann, V. Sentchilo, C. von Mering, J. R. van der Meer, Reproducible propagation of species-rich soil bacterial communities suggests robust underlying deterministic principles of community formation. *mSystems*, e0016022 (2022).10.1128/msystems.00160-22
